# Transcriptomic correlates of electrophysiological and morphological diversity within and across neuron types

**DOI:** 10.1101/524561

**Authors:** Claire Bomkamp, Shreejoy J. Tripathy, Carolina Bengtsson Gonzales, Jens Hjerling-Leffler, Ann Marie Craig, Paul Pavlidis

## Abstract

In order to further our understanding of how gene expression contributes to key functional properties of neurons, we combined publicly accessible gene expression, electrophysiology, and morphology measurements to identify cross-cell type correlations between these data modalities. Building on our previous work using a similar approach, we distinguished between correlations which were “class-driven,” meaning those that could be explained by differences between excitatory and inhibitory cell classes, and those that reflected graded phenotypic differences within classes. Taking cell class identity into account increased the degree to which our results replicated in an independent dataset as well as their correspondence with known modes of ion channel function based on the literature. We also found a smaller set of genes whose relationships to electrophysiological or morphological properties appear to be specific to either excitatory or inhibitory cell types. Next, using data from Patch-seq experiments, allowing simultaneous single-cell characterization of gene expression and electrophysiology, we found that some of the gene-property correlations observed across cell types were further predictive of within-cell type heterogeneity. In summary, we have identified a number of relationships between gene expression, electrophysiology, and morphology that provide testable hypotheses for future studies.

**Author Summary:** The behavior of neurons is governed by their electrical properties, for example how readily they respond to a stimulus or at what rate they are able to send signals. Additionally, neurons come in different shapes and sizes, and their shape defines how they can form connections with specific partners and thus function within the complete circuit. We know that these properties are governed by genes, acting acutely or during development, but we do not know which specific genes underlie many of these properties. Understanding how gene expression changes the properties of neurons will help in advancing our overall understanding of how neurons, and ultimately brains, function. This can in turn help to identify potential treatments for brain-related diseases. In this work, we aimed to identify genes whose expression showed a relationship with the electrical properties and shape measurements of different types of neurons. While our analysis does not identify causal relationships, our findings provide testable predictions for future research.

## Introduction

Two prominent features that distinguish neurons from other cells are their electrical activity and their characteristic morphology. The specific pattern of electrophysiological activity displayed by a given neuron is a core property of its identity as one type of neuron or another. Similarly, different cell types often show striking differences in their size, branching complexity, and other morphological features. Neuronal cell types defined according to their electrophysiological or morphological characteristics show substantial correspondence with one another as well as with those defined using classification schemes based on transcriptomic criteria (1). Electrophysiological characteristics of neurons, as well as their connectivity patterns, give rise to the computational properties of a given circuit (2,3). Additionally, modeling studies show that morphological changes in simulated neurons can critically change their signaling capabilities (4–6). Thus, understanding the origins of neuronal electrophysiology and morphology is an important step in understanding the mechanisms of brain function, both in the context of basic research and in the search for treatments for neuropsychiatric disorders.

A comprehensive understanding of the mechanisms that give rise to electrophysiological or morphological diversity must necessarily include a catalogue of the genes whose products contribute to these properties. Many genes have been shown experimentally to influence neuronal electrophysiology through a variety of mechanisms, including but not limited to ion channel activity, protein trafficking, and transcription factor activity (7–9). Processes such as axon guidance and the development of dendrite morphology are also known to be under genetic control (10). Despite this, our understanding of the relationship between gene expression and electrophysiological or morphological properties is quite limited.

In previous work (11), we combined publicly accessible electrophysiological and gene expression datasets in order to examine the relationship between gene expression and electrophysiological properties. By matching groups of cells inferred to be similar based on multiple information sources, such as the transgenic reporter line and the brain region cells were isolated from, we were able to combine separate datasets containing gene expression and electrophysiological data to generate lists of genes which were correlated with one of several electrophysiological properties (as outlined in Fig 1A). The goal of this approach was to identify candidate genes that could be further studied using knockout or knockdown approaches in order to determine whether a causal relationship was present.

**Fig 1.**
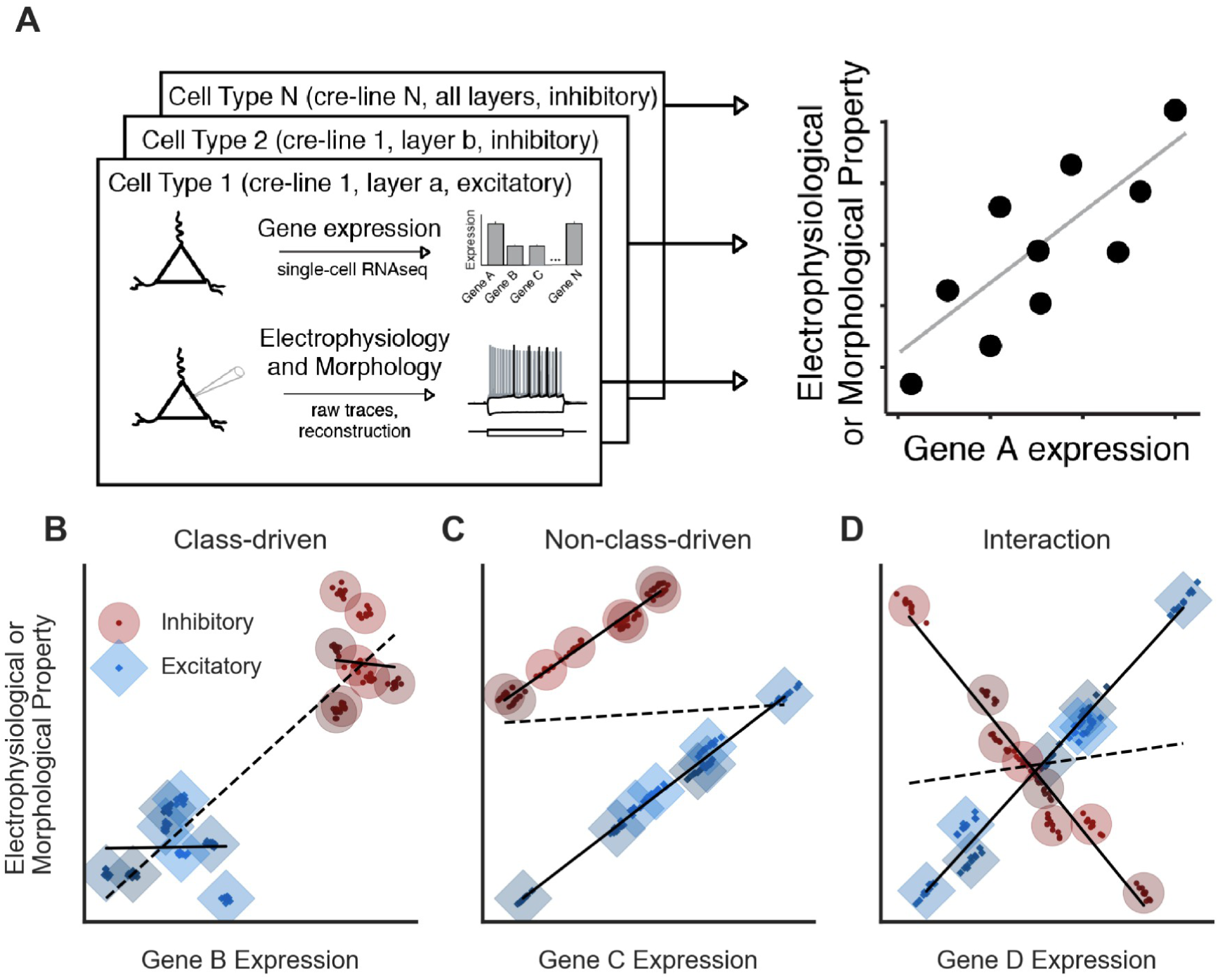
Methods for modeling relationships between gene expression and electrophysiological or morphological properties with respect to cell class. A. Schematic for defining cell types from single-cell transcriptomic or electrophysiological and morphological data. We divided cells into types based on Cre-driver expression as well as cortical layer and excitatory/inhibitory identity (left). Right panel shows summarization of cellular features by cell type for a hypothetical gene and property, where each point in the scatter plot represents each cell type’s mean gene expression (x-axis) and the mean value of an electrophysiological or morphological property (y-axis). B. A hypothetical class-driven relationship between a gene and an electrophysiological or morphological property, in which neither cell class (excitatory or inhibitory) shows a relationship between gene expression and the property (solid lines), but an overall relationship appears because of systematic cross-class differences in both data modalities (dashed line). For B-D, small points represent individual cells and larger circles or diamonds represent cell type averages. C. A hypothetical example of a non-class-driven relationship, where the gene-property relationship appears within each major cell class (solid lines), but would be obscured if modeled in a class-independent manner (dashed line). D. A hypothetical example of a gene-property relationship exhibiting an interaction with cell class. Here, expression of the gene is positively correlated with the property in excitatory cell types but negatively correlated in inhibitory types (solid lines).

One caveat in our prior study is that the gene-electrophysiology correlations we identified may have been confounded by overall differences between broad cell classes. Across multiple datasets and cellular characterization methods, including gene expression (12–15), and electrophysiology and morphology (1), clustering cellular phenotypes in an unbiased manner reveals the major taxonomic difference between neurons to be between projecting and non-projecting neurons (13), or in the case of those cell types present in the cortex or hippocampus, excitatory and inhibitory neurons (12,14,15). Thus, the commonly held view that a neuron’s identity is first and foremost defined by its excitatory or inhibitory identity is corroborated across multiple data sources and experimental modalities.

Therefore, we reasoned that the dataset we used previously was potentially susceptible to this confounding effect of cell class, since it contained a mixture of cells from different broad cell classes. In this work, we will use the term “cell type” to refer to narrowly-defined cell types, and “cell class” to refer to those which are broadly-defined (excitatory versus inhibitory or projecting versus non-projecting). We refer to correlations between gene expression and electrophysiological or morphological properties that are explained by differences between cell classes as “class-driven,” (e.g. Fig 1B) and to those that exist based on graded differences within broad cell classes as “non-class-driven” (e.g. Fig 1C). We reason that gene-property relationships that are non-class-driven would be more likely to be potential causal regulators of the associated property. Although some class-driven correlations likely do reflect true relationships between genes and properties which distinguish excitatory from inhibitory cells, separating these relationships from instances where one cell class has a higher value of a property and coincidentally higher or lower expression of a gene without additional sources of data is not possible. Effectively, such situations are analogous to attempting to draw conclusions about correlations with only two data points.

Due to limitations in available data, we were unable to address the effect of cell class in our previous work (11). Since then, the RNA-seq and electrophysiology datasets from the Allen Institute for Brain Science (AIBS) (which we originally used as validation data) have expanded greatly, with more cells and more transgenic lines represented. This increase in size, together with the fact that the AIBS data were collected using standardized protocols, suggests that this dataset might prove valuable for discovering genes correlated with electrophysiological and morphological properties. In addition, the growing use of the Patch-seq methodology (17), allowing transcriptomic, electrophysiological, and morphological characterization of the same single cell, also affords an opportunity to test gene-property correlations.

Leveraging the larger size of the new AIBS dataset, we were able to address limitations of our previous study related to excitatory versus inhibitory cell class by employing statistical methods to help mitigate the effects of cell class. These methods, together with the larger number of cell types represented in the new dataset, allowed us to identify novel electrophysiological and morphological property-related gene sets which are potentially more likely to represent meaningful biological relationships.

## Results

### Primary Dataset

The primary dataset we used combined groups of cells from mouse visual cortex characterized by the Allen Institute for Brain Science (AIBS; http://celltypes.brain-map.org/), where multiple Cre-driver lines were used to target cells for characterization. Standard electrophysiological protocols were used to characterize cells *in vitro*, with a subset of these cells further undergoing detailed morphological characterization (1). In addition, a separate group of cells were subjected to deep single-cell RNA-sequencing to characterize cellular transcriptomes (14). Because the same Cre-lines were used to characterize cells along multiple modalities of neuronal function, we were able to summarize these data to the “cell type” level (reflecting Cre-line, cortical layer, and major neurotransmitter; shown in Table S1) by pooling and combining cellular characterization data across different animals and data modalities. The definition of multiple cell types within one Cre-line based on cortical layer and major neurotransmitter is supported by cross-layer differences in gene expression (14) and in electrophysiological properties (Fig S1).

The final combined dataset is composed of 34 inhibitory GABAergic and 14 excitatory glutamatergic types (48 total) with electrophysiological data, and 30 inhibitory and 13 excitatory types (43 total) with morphological data. The increased size of this dataset is a considerable advance over our prior analysis (11), which employed an older version of the same dataset (only 12 cell types) (15). This was made possible in part because of more Cre-lines available for analysis and finer cortical layer dissections for the transcriptomic data. For each cell type thus defined, we computed the mean expression value for each gene represented in the RNA-seq dataset and the mean value of each of sixteen electrophysiological and six morphological properties (described in Table S2).

### Analysis Approach

Our goal was to identify, for each electrophysiological or morphological property, genes that were correlated with the property (Fig 1A). However, overall differences between excitatory and inhibitory cell classes can make the interpretation of such relationships more complicated in several ways. For example, Fig 1B shows an example of a gene-property correlation that appears almost entirely **class-driven**, meaning that although no relationship appears *within* either cell class, the apparent relationship is entirely driven by differences *between* cell classes. In this case, inhibitory cell types show higher expression of the gene and a greater value of the property compared to excitatory cell types. In contrast, Fig 1C shows a **non-class-driven** relationship, meaning one that manifests in both cell classes, but which may be obscured by baseline differences when the cell classes are grouped. In this example, a correlation that appears within both classes independently is obscured by a higher value of the property in inhibitory compared to excitatory cell types. Although this obscuring effect is present in this particular example, it is not required for a relationship to be considered non-class-driven; we expected to see some relationships that were consistent both within each class as well as among all cell types.

In order to computationally account for these possibilities, we evaluated each combination of gene and property using a statistical model that assesses the predictive value of the gene on the property while controlling for the effects of cell class. We termed this model the **class-conditional model**. This model would be expected to identify a significant result when a non-class-driven relationship is present (Fig 1C), but would not identify relationships that are class-driven (Fig 1B). For comparison, we modeled the same gene-property pairs using a **class-independent model**, which assesses the predictive value of the gene on the property irrespective of cell class. This model is similar in principle to the correlational method used in our previous work (11) and would be expected to produce a significant result in cases showing class-driven relationships (such as Fig 1B) but might miss some instances of non-class-driven relationships (such as Fig 1C).

Another possible gene-property relationship is one where there is an interaction between gene and class, meaning that the gene-property relationship is different in excitatory and inhibitory cell types. An interaction could indicate either that excitatory and inhibitory cell types both show a correlation between the gene and property, but the slopes are in opposite directions (as in the example in Fig 1D), or that the gene is correlated with the property only in one cell class. To detect such situations, we introduced a third model, the **interaction model**, which tested whether the relationship between gene expression and the property in question was significantly different between excitatory and inhibitory cell types. In summary, the three models are designed to answer three different questions:

Class-independent model: Is expression of the gene a significant predictor of the property if we assume that cell class is not a factor?

Class-conditional model: After accounting for cell class, is the gene’s expression a significant predictor of the property?

Interaction model: Is the relationship between the gene’s expression and the property statistically different in inhibitory and excitatory cells?

### Accounting for cell class results in the identification of a distinct but overlapping set of genes

We first set out to understand how accounting for cell class identity (excitatory or inhibitory) affects the interpretation of gene-property relationships. We modeled each relationship with or without including an indicator variable for cell class, using the class-conditional or class-independent models described above. For most properties, we found that the degree of overlap between the sets of genes identified in the two models (at a false discovery rate (FDR) < 0.1) was substantial but far from a complete intersection (Fig 2A, Venn diagrams, and Table S2). For example, for after-hyperpolarization (AHP) amplitude, we found ~6000 significantly-associated genes in the class-independent model and ~6500 in the class-conditional model; out of these, ~3700 genes were shared between models. Thus, accounting for cell class results in the identification of a substantially different set of candidate genes, which suggests that many of the genes identified in our previous work (11) might reflect class-driven gene-property relationships.

**Fig 2.**
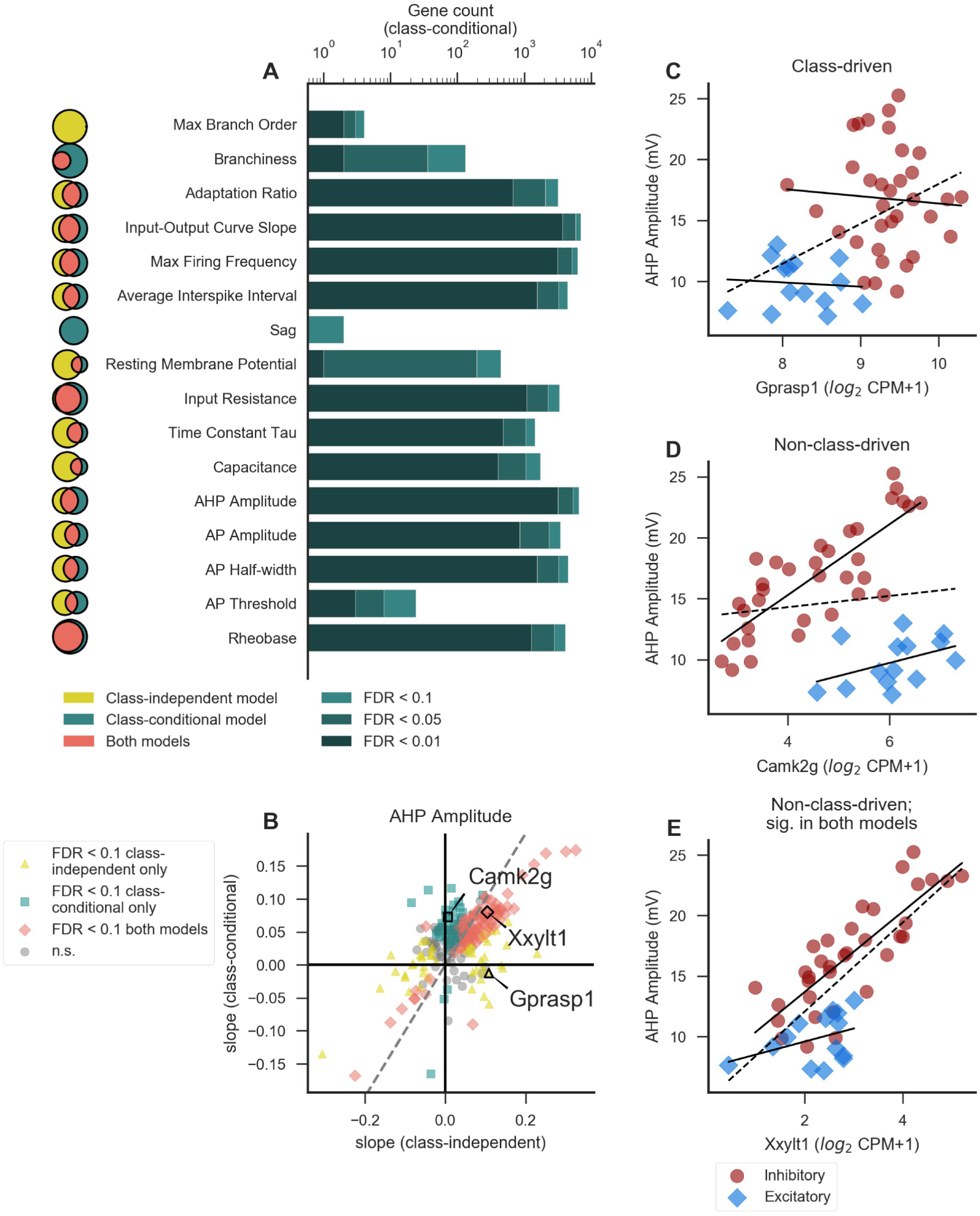
Different sets of genes are associated with electrophysiological and morphological properties after correcting for cell class. A. Number of genes significantly associated with each property in the class-conditional model at various levels of significance (only properties with significant genes in this model are shown). Darkness of the bar represents the significance level of each group of genes. Venn diagrams to the left indicate the extent of overlap (pink; middle) between the gene sets identified by the class-independent (gold; left) and class-conditional (teal; right) models, where the area of each segment is proportional to the significant gene count at a threshold of FDR < 0.1. Venn diagrams for different properties are not to scale with one another. See Table S2 for descriptions of electrophysiological and morphological properties analyzed here, as well as gene counts for all properties. B. Comparison of model-based slopes from the class-independent and class-conditional models. Each point represents a single gene’s relationship with the electrophysiological property AHP amplitude and is colored according to whether the relationship is significant in one or both models (FDR < 0.1). Example genes in C-E are indicated. For clarity of visualization, only a random subset of genes (2% total) are shown to mitigate over-plotting‥ Dashed line indicates identity. C-E. Examples of genes showing significant associations with AHP amplitude that are class-driven (C; significant in class-independent model only), non-class-driven (D; significant in class-conditional model only), or non-class-driven but significant by either model (E). Solid lines indicate linear fits within excitatory or inhibitory cell classes only and dashed line indicates a linear fit including all cell types. Gene expression is quantified as counts per million (CPM).

We next asked how overall differences in morphological and electrophysiological properties between excitatory and inhibitory cells affect gene-property relationships. To this end, we used a linear model to estimate the effect of cell class on each property. For most properties, there was a significant (p < 0.05) effect of cell class. The features of action potential (AP) threshold, input resistance, sag, rheobase, branchiness, soma surface, and bifurcation angle are exceptions to this. The existence of a significant difference in most properties between excitatory and inhibitory cell types highlights the importance of taking cell class into account when attempting to relate these properties to gene expression. The properties without a significant difference are likely to be less susceptible to class-driven effects, but the class-independent model still might miss potentially interesting relationships due to differences in gene expression between classes, resulting in genes which are identified by the class-conditional model only.

We compared the strength and direction of the relationship in both the class-independent and class-conditional models by directly comparing the slopes derived from each model for each gene-property relationship (where slope indicates the change in the property per 2-fold change in gene expression; shown for AHP amplitude in Fig 2B). While there is broad agreement between the class-independent and class-conditional models (r_Spearman_ = 0.52), a substantial number of gene-property relationships are significant in one model but not the other (FDR < 0.1). In other words, these relationships are either class-driven (significant in the class-independent model only) or non-class-driven and obscured by class (significant in the class-conditional model only). For example, the relationship between the gene *Gprasp1* and AHP amplitude illustrates an example of a class-driven relationship where the apparent relationship is entirely due to broad differences in excitatory and inhibitory classes (Fig 2C). The gene *Camk2g* shows a non-class-driven relationship with the same property that is obscured in the class-independent model by higher AHP amplitude values in inhibitory cell types (Fig 2D). However, many genes, such as *Xxylt1*, are identified using either model (Fig 2E).

### Divergent gene-property relationships in inhibitory versus excitatory cell classes

We next wondered whether some gene-property relationships might be potentially different within, or specific to, excitatory or inhibitory cell types. To test this, we incorporated an interaction term between gene expression and excitatory versus inhibitory cell class to assess whether the gene-property relationships (i.e. slopes) were different within each cell class. For nearly all properties, there were fewer significant genes in the interaction model compared to the class-conditional model (Fig 3A, Venn diagrams, and Table S3). For example, out of the ~6500 genes significantly associated with AHP amplitude in the class-conditional model, ~2000 also show interactions, and there are an additional ~700 which show an interaction but are not significant in the class-conditional model. This could indicate that “true” interactions are comparatively rare, but this finding is also likely partly explained by differences in statistical power. In addition, these interactions do not appear to be merely the result of low or no gene expression within one cell class but not the other; we did not observe strong correlations for any property between the interaction model slope and the average difference in expression levels between inhibitory and excitatory cell types (Fig S2).

**Fig 3.**
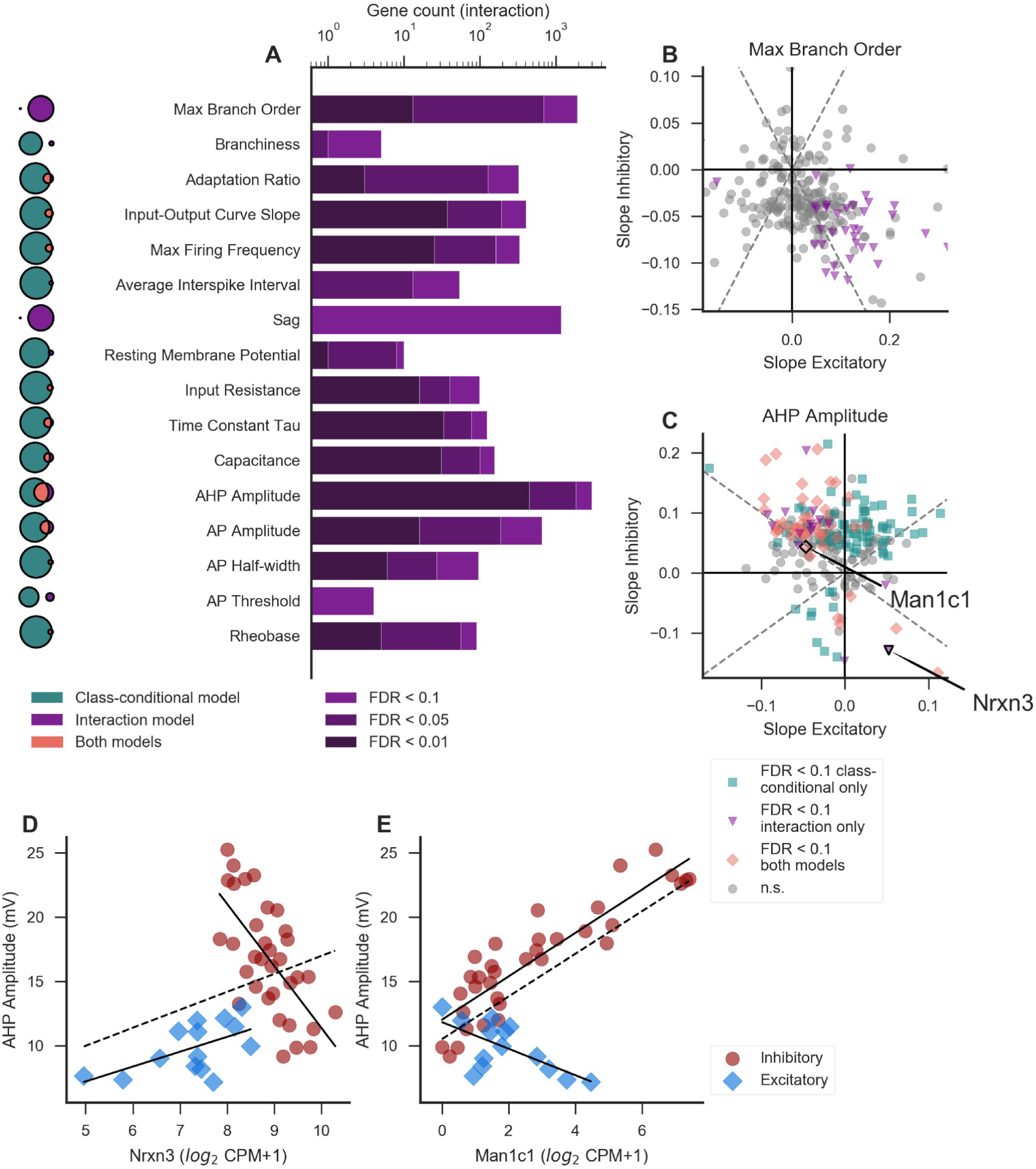
Identification of divergent gene-property relationships in excitatory versus inhibitory cell classes. A. Number of genes showing a significant interaction effect between gene and class for each property. Darkness of the bar represents the significance level of each group of genes. Venn diagrams to the left indicate the extent of overlap (pink; middle) between the class-conditional (teal; left) and interaction (purple; right) models, where the area of each segment is proportional to the significant gene count at a threshold of FDR < 0.1. Venn diagrams for different properties are not to scale with one another. B-C. Slope values within excitatory cell types (x axis) plotted against the slope values for the same set of genes in inhibitory cell types (y axis). Each point represents a single gene’s relationship to the morphological property maximum branch order (B) or electrophysiological property AHP amplitude (C), and is colored according to its significance in one or both models (see inset legend). Example gene-property relationships highlighted in D-E are marked in panel C. For clarity of visualization, only a random 2% subset of the total number of genes are plotted. Dashed lines indicate positive and negative unity lines. D. Example of a gene with a significant interaction term which is not significant in the class-conditional model. For D and E, solid lines indicate linear fits including only excitatory or only inhibitory cell types, and dashed line indicates a linear fit including all cell types. E. Example of a gene which is significant in both the class-conditional and interaction models.

For all properties, we found that the slopes of the gene-property relationships within excitatory cell types were poorly correlated with those within inhibitory cell types (example features maximum branch order and AHP amplitude shown in Fig 3B, C). By definition, the genes with significant interaction terms were those where the slopes calculated within excitatory and inhibitory classes were very different from each other (pink and purple points in Fig 3B, C). If the majority of gene-property relationships are shared between excitatory and inhibitory cell types, as suggested by the greater number of significant genes in the class-conditional model than in the interaction model for most properties, one might expect a positive correlation between slopes calculated in inhibitory and excitatory cell types. However, such a correlation may be lacking in this analysis because we would expect most genes to have no relationship to a given property and thus most slopes to be near zero.

The properties maximum branch order and sag are unusual in that they show few significant genes using the class-conditional model, but many (1914 and 1174, respectively) in the interaction model (Fig 3A, Venn diagrams, and Table S3; slopes for maximum branch order plotted in Fig 3B). We hypothesize that this might be because these properties are under stronger (or otherwise more readily identified) genetic control in excitatory compared to inhibitory cell types (see Discussion).

Fig 3D, E show examples of genes with significant interaction terms for AHP amplitude. The class-conditional model also shows a significant relationship in the case of *Man1c1* (Fig 3E) but not *Nrxn3* (Fig 3D). In other words, the interaction model identified a potentially interesting relationship in the case of *Nrxn3* which was missed by the class-conditional model. For *Man1c1*, the interaction model does not reveal a new relationship, but instead highlights the fact that this gene-property relationship, if real, is potentially more complicated than would be assumed based on the class-conditional model alone. *Man1c1* is an enzyme involved in the maturation of N-linked oligosaccharides (18), and is thus a plausible regulator of AHP amplitude, since N-linked glycosylation of voltage-gated potassium channels or their auxiliary subunits is known to regulate both surface trafficking and channel function (19,20). The apparent class-specificity of this relationship could result from class-specific co-expression of certain potassium channels or other enzymes involved in glycan synthesis or maturation.

### Results from the class-conditional model are more likely to validate using independent methods

We next asked how the gene-property relationships from the class-independent and class-conditional models, based on our analysis of the AIBS cortical cell types dataset, might generalize to other datasets. We first compared the results reported here to those from our earlier NeuroElectro/NeuroExpresso (NE) literature-based dataset (11), after subsetting these data to include only non-projecting cell types (reflecting 19 cell types in total sampled throughout the brain, described in detail in the Methods). We chose to use non-projecting cell types in the NE dataset, as these were recently described by a mouse brain-wide transcriptomic survey as corresponding to a single broad cell class (13). To this end, we calculated Spearman correlations between genes and electrophysiological properties in the NE dataset. Next, for gene-property relationships from both the class-independent and class-conditional models, we assessed their aggregate consistency with those from the NE dataset. Here, we defined “consistency” for a given model (i.e. class-independent or class-conditional) and property as the correlation between gene-property slopes calculated from the AIBS dataset with the Spearman correlations for the same set of gene-property relationships in the NE dataset (illustrated in Fig 4B).

**Fig 4.**
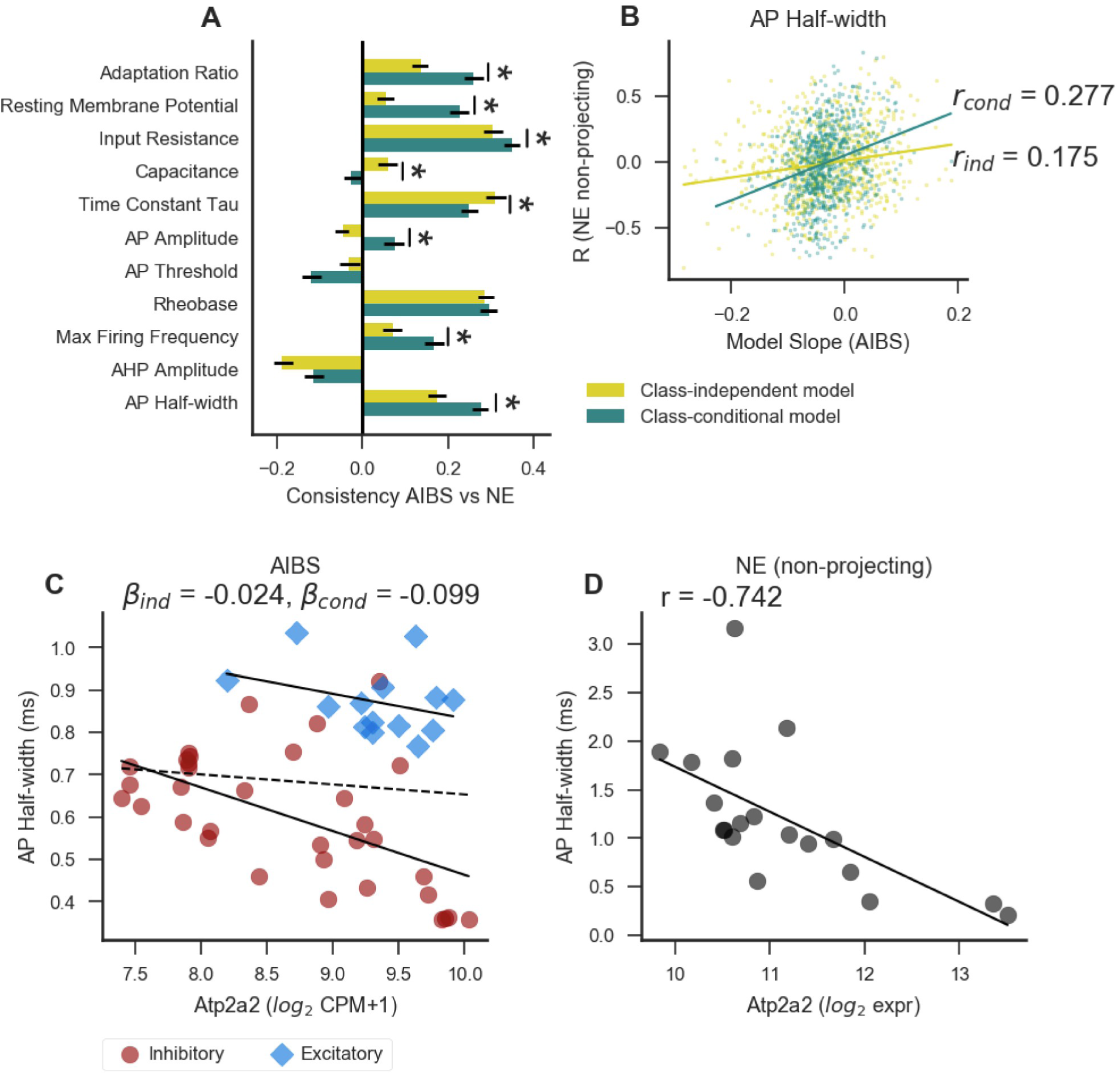
Modeling genelelectrophysiology relationships using the class-conditional model is more predictive than the class-independent model of correlations in an independent dataset containing non-projecting cell types only. A. Aggregate gene-property relationship consistency between AIBS and NeuroExpresso /NeuroEle ctro (NE) datasets. Error bars indicate a 95% confidence interval, and asterisk indicates a significant (p < 0.05) difference in the consistency metric between the class-independent and class-conditional models, calculated using 100 bootstrap resamples of the original values (not indicated for properties where both values are negative). B. Direct comparison of gene-property relationships between the AIBS and NE datasets. Each point represents the relationship between a single gene and the property AP half-width. The model slope from the AIBS dataset is plotted on the x axis (with the class-independent model (ind) slopes in gold, and the class-conditional model (cond) slopes in teal), and the Spearman correlation for the same set of genes in the NE dataset on the y axis. For clarity of visualization only 10% of the total number of genes are plotted. Lines indicate a linear fit for each set of points. The correlation within each set of points is used as a measure of cross-dataset consistency (plotted for all properties in panel A). C-D. Example of a gene showing consistent results between the NE dataset and the AIBS dataset using the class-conditional model, but not the class-independent model. C shows the relationship within the AIBS dataset, and D shows the same gene and property in the NE dataset. Solid lines indicate a linear fit including only types belonging to one cell class, and dashed line indicates a linear fit including all cell types.

In Fig 4A we show a comparison of the gene/electrophysiology correlations from the AIBS dataset with the model slopes (beta) from the NE dataset (11). We found that for seven out of the eleven electrophysiological properties shared between the datasets, both AIBS dataset-based statistical models were consistent with analogous gene-property relationships based on the NE dataset (r_Spearman_ as high as 0.305 and 0.35 for class-independent and class-conditional, respectively). For six out of the eleven features, we found that the class-conditional model was considerably more consistent than the class-independent model with relationships in the NE dataset. For only two features, capacitance and membrane time constant (tau), was the class-independent model more consistent than the class-conditional with the NE dataset. Fig 4B shows an example of how consistency was measured for AP half-width. The relationship between *Atp2a2* expression and AP half-width is shown in Fig 4C, D as an example of a gene-property relationship which is consistent between the NE (r = −0.742) and AIBS datasets for the class-conditional (beta = −0.099 ± 0.024; FDR = 0.002) but not the class-independent (beta = −0.024 ± 0.034; FDR = 0.62) model.

### Assessing within-cell type correlations using Patch-seq datasets

We next wondered whether these between-cell type gene-property relationships might be predictive of cell-to-cell heterogeneity within a given cell type. We reasoned that the recently developed Patch-seq methodology, allowing morphological, electrophysiological, and transcriptomic characterization from the same single cell, presents a unique opportunity to test this possibility (17). While these data at present are limited by relatively modest sample sizes and technical factors such as inefficient mRNA capture and potential off-target cellular mRNA contamination (21), we nonetheless sought to use these data to assess the nature of within-cell type gene-property relationships.

To this end, we performed an integrated analysis of 5 Patch-seq datasets, with each dataset characterizing transcriptomic and electrophysiological diversity of mouse forebrain inhibitory cells from the neocortex, hippocampus, and striatum (Table 1). Our analysis includes one novel dataset of 19 Pvalb-Cre positive interneurons recorded in region CA1 of the mouse hippocampus, reported here for the first time. Cells in this dataset (referred to as the Bengtsson Gonzales dataset), were characterized as described in (22).

**Table 1.** Description of Patch-seq datasets re-analyzed in this study. Depending on the dataset, RNA amplification was performed using variations on single-cell-tagged reverse transcription (STRT) (28) or Switching Mechanism At the end of the 5’-end of the RNA Transcript (SMART) (29). The Bengtsson Gonzales dataset reflects a novel dataset reported here for the first time.

| Dataset | Description | RNA amplification | Number of cells | Accession |
| --- | --- | --- | --- | --- |
| Cadwell (17) | Cortical layer 1 interneurons | Smart-seq2 | 57 | E-MTAB-4092 |
| Fuzik (26) | Cortical layer 1/2 interneurons and pyramidal cells | STRT-C1 (with unique molecule identifiers) | 80 | GSE70844 |
| Földy (27) | Hippocampal CA1 and subiculum pyramidal cells and regular- and fast-spiking interneurons | SMARTer | 93 | GSE75386 |
| Muñoz-Manchado (22) | Striatum interneurons | STRT-C1 (with unique molecule identifiers) | 99 | GSE119248 |
| Bengtsson Gonzales | Hippocampal CA1 Pvalb-Cre interneurons | STRT-C1 (with unique molecule identifiers) | 19 | N/A |

To jointly analyze these Patch-seq datasets, we first mapped Patch-seq sampled cells to the *cell type level*, using a transcriptome-based classifier that assigns cells to cell types as defined by cellular dissociation-based single-cell RNAseq reference atlases from the cortex and striatum (14,22). Specifically, we resolved individual cells to the level of major cell types; for example, Pvalb, Sst, Vip, Lamp5, etc. (referred to in Tasic et al., 2018 as “subclasses”). Next, for each cell type, we identified genes that are highly variable in their expression levels *within* cells of the same type. We reasoned that these highly-variable genes might be those most likely to drive or appear correlated with electrophysiological heterogeneity within each cell type. Lastly, we performed a joint analysis across Patch-seq datasets to assess the strength of gene-property relationships within cell types where the gene was highly variable. Here, we used a mixed-effects regression model, with gene expression as a fixed effect and dataset and cell type as random effects and with cells weighted by their estimated transcriptome quality (see Methods).

Despite the limitations of the Patch-seq data, we found a small number of genes whose expression levels were significantly associated with cell-to-cell electrophysiological heterogeneity within cell types (FDR < 0.1; Fig 5A). For example, we found that expression of *Kcna1*, which encodes the potassium channel Kv1.1, was inversely correlated with AP half-width (Fig 5B; Beta_Patch-seq_ = −0.0484 ± 0.0106, FDR_Patch-seq_ = 0.0683) within hippocampal *Pvalb* and striatum *Pthlh* cells (the only cell types in which the variability in *Kcna1* expression met our threshold for analysis). Importantly, there was also a significant relationship with the same directionality for *Kcna1* and AP half-width in the AIBS dataset (Beta_class-conditional_ = −0.048 ± 0.011, FDR_class-conditional_ = 0.001). Moreover, the relationship between *Kcna1*/Kv1.1 expression and action potential width has been experimentally reported previously (23) (Brew et al., 2003).

**Fig 5.**
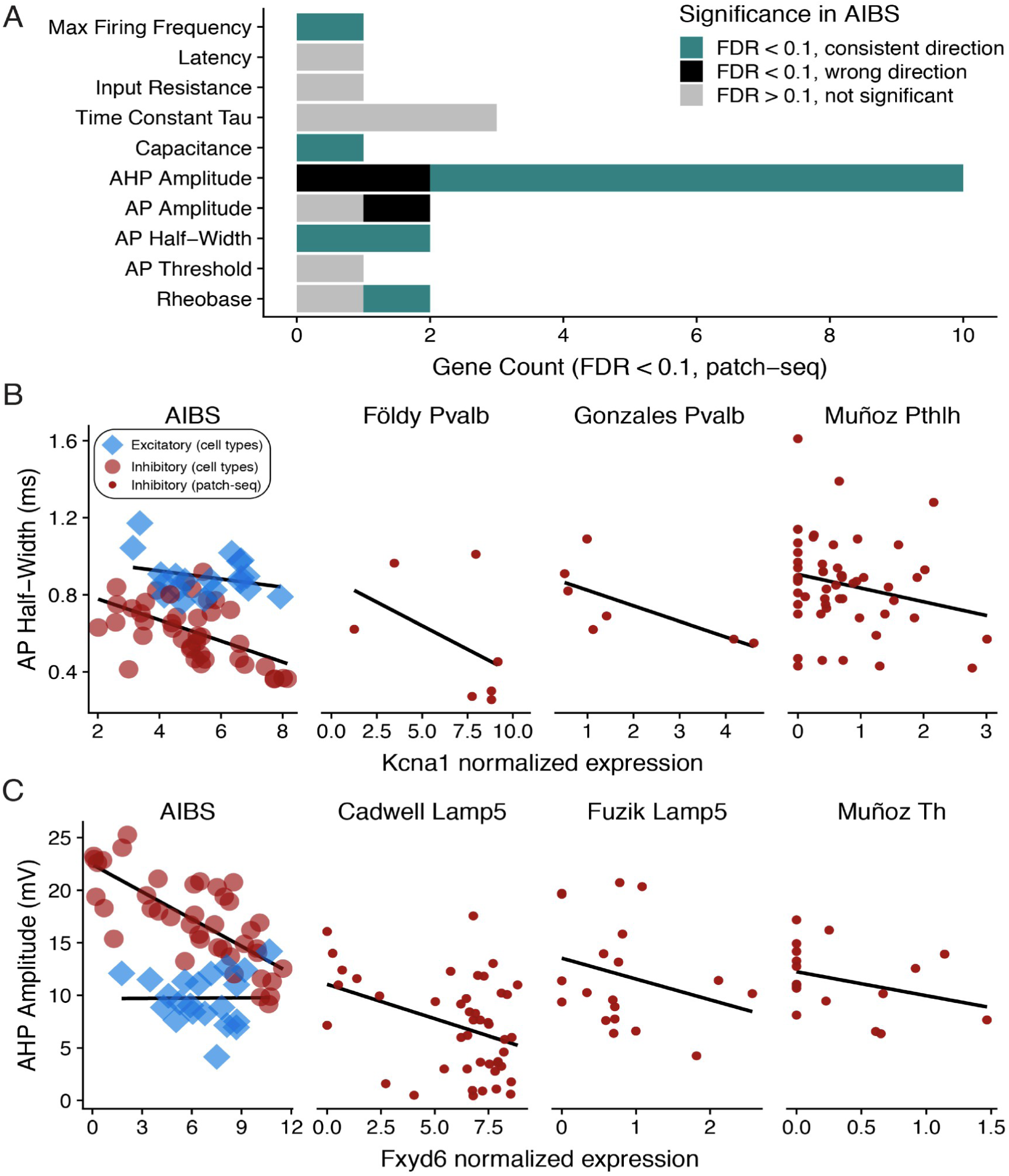
Assessing gene-property relationships within cell subclasses using Patch-seq. A. Number of genes associated with each electrophysiological property based on a joint cross - laboratory analysis of 5 Patch-seq datasets. Genes shown are significant at FDR < 0.1, based on a mixed-effects regression model, treating gene expression as a fixed effect and dataset identity and cell type as random e ffects. Bar color denotes overlap of Patch-seq based gene-property relationships with analogous relationships from the AIBS class-conditional model analysis. Note that analysis of gene - property relationships in the Patch-seq datasets are independent from those in the AIBS cell types analysis. B, C. Examples of genes showing significant associations with electrophysiological features in the class-conditional analysis of the AIBS dataset (left-most panel) and the mixed-effects analysis of the Patch-seq datasets (other panels). Dataset name and cell type is shown in the subpanel title and solid lines indicate linear fits within cell classes (AIBS) or fits within each Patch-seq dataset and cell type, after weighting cells by transcriptome-quality (see Methods). Based on differences in mRNA quantification, x-axis units for AIBS, Cadwell, and Földy datasets are log2 (CPM+1), and for Bengtsson Gonzales, Muñoz, and Fuzik datasets are log2 normalized molecule counts (normalized to 2000 unique molecules per cell).

As another example, we saw an inverse correlation between *Fxyd6* expression and AHP amplitude, based on cortical Lamp5- and striatum Th- cells (Fig 5C, Beta_Patch-seq_ = −0.695 ± 0.118, FDR_Patch-seq_ = 0.00841). We also saw a similar relationship in the AIBS dataset (Beta_class-conditional_ = −0.021 ± 0.003, FDR_class-conditional_ = 0.00001). Intriguingly, *Fxyd6* encodes phosphohippolin, a regulator of Na+/K+ ATPase activity (24) and is thus plausibly involved in the AHP and action potential repolarization. Intriguingly, in a separate single-cell RNA-seq study of CA1 interneurons, *Fxyd6* was found to be more highly expressed cells known to spike more slowly (25).

In general, we found that when a gene-property relationship was statistically significant in both the Patch-seq and AIBS class-conditional analyses (FDR < 0.1), this relationship was usually in the same direction in both analyses (Fig 5A; 10 out of 13 gene-property relationships total). Results were similar in the class-independent model, except with a smaller set of gene/ephys relationships matching between both (7 out of 9 relationships were in a consistent direction). All of the genes which were consistent between the class-independent and Patch-seq analyses were also consistent in the class-conditional model. While our analyses of these Patch-seq datasets should be considered preliminary (pending the availability of larger and higher-quality datasets), we find the correspondence with our earlier analysis encouraging. Namely, this analysis suggests that some of the same genes that appear to drive large differences across cortical cell types might also be defining more subtle within-cell type heterogeneity.

### The expected relationship between voltage-gated potassium channels and AHP amplitude is apparent only after accounting for cell class

We next asked whether we see a relationship between an electrophysiological feature and a category of genes which are known regulators of that feature. Voltage-gated potassium channels are known to be involved in producing the after-hyperpolarization following an action potential (30,31) (AHP amplitude; illustrated by the dashed arrow in Fig 6A). We thus hypothesized that for many of these genes, higher expression levels would be associated with larger AHP amplitudes (although not all voltage-gated potassium channels necessarily contribute directly to AHP amplitude). We further hypothesized that this relationship would be more apparent after accounting for cell class, in part because AHP amplitudes differ considerably between excitatory and inhibitory cell classes (Fig 6B-D). Indeed, our previous work found a spurious negative correlation between expression of the *Kcnb1* gene and AHP amplitude which resulted from higher expression of Kcnb1 in excitatory cell types compared to others (11).

**Fig 6.**
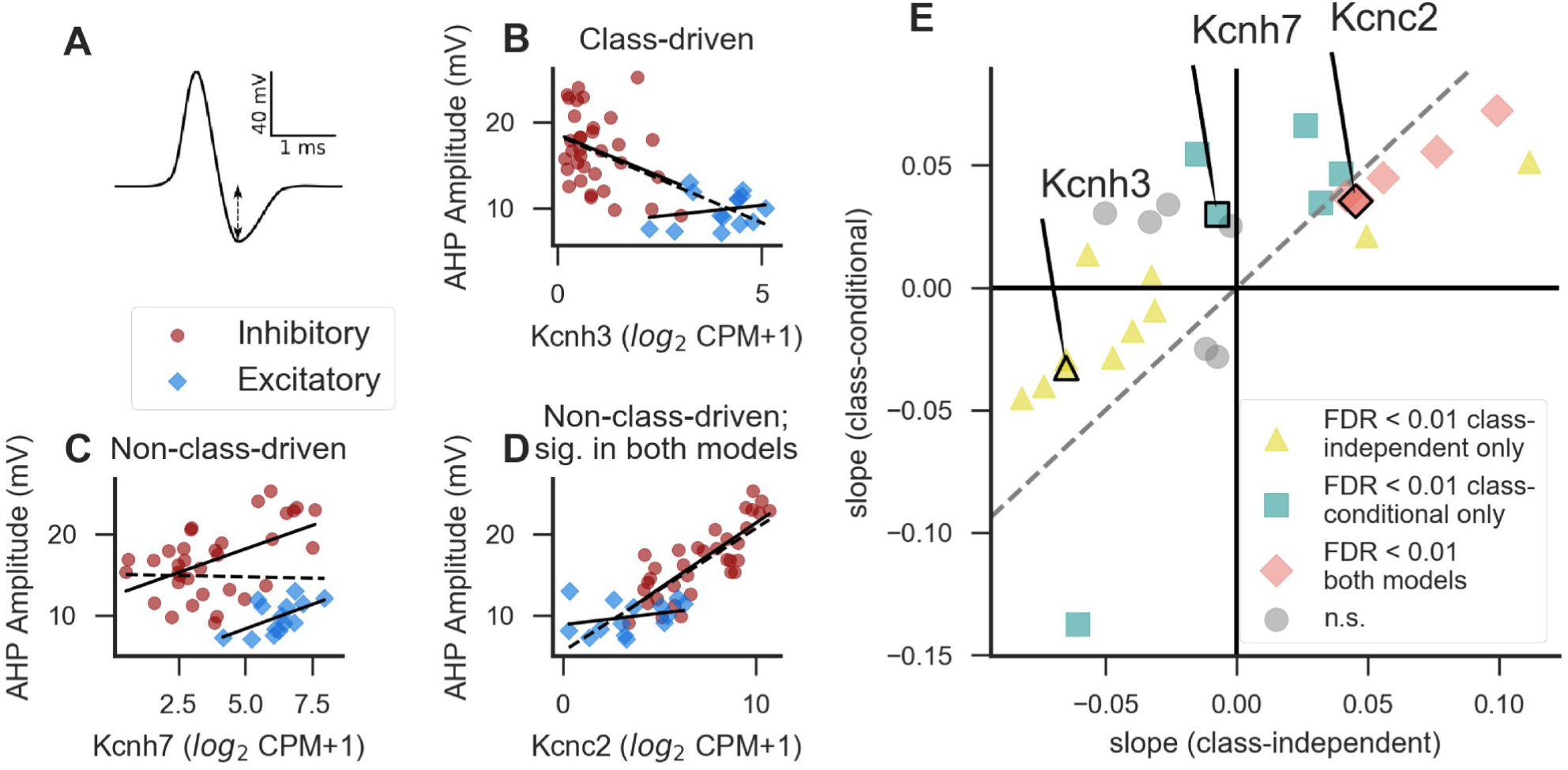
Accounting for cell class changes the interpretation of the relationship between potassium channel expression and after-hyperpolarization amplitude. A. Schematic view of an action potential trace, with the dashed line representing the ARP amplitude value. B-D. Examples of voltage-gated potassium channel genes significantly associated with ARP amplitude in the class-independent model (B), the class-conditional model (C), or both (D) at a threshold of FDR < 0.01. Solid lines indicate a linear fit including only excitatory or only inhibitory cell types, and dashed line indicates a linear fit including all cell types. E. Comparison of class-independent and class-conditional approaches for detecting associations between voltage-gated potassium channels and ARP amplitude. Each point indicates a single gene, and x and y axes are the slopes from the class-independent and class-conditional models, respectively. Labeled points are the example genes shown in B-D. Dashed line indicates identity.

We evaluated model slopes between each of 29 voltage-gated potassium channel genes (32) and AHP amplitude in the AIBS dataset for each of the class-independent and class-conditional statistical models (examples shown in Fig 6B-D and summary in Fig 6E).

Examples of voltage-gated potassium channel genes associated with AHP amplitude include *Kcnh3* (Fig 6B) in a class-driven and *Kcnh7* and *Kcnc2* in a non-class-driven manner (Fig 6C, D). In total, the class-independent model identified 17 significant genes (at a stringent threshold of FDR < 0.01), with 8 of these genes having positive slopes and 9 negative. In contrast, there were 12 genes that were significantly associated with AHP amplitude in the class-conditional model at the same statistical threshold, and 11 of these genes had slopes in the positive direction. Thus the results obtained using the class-conditional model are consistent with our *a priori* hypothesis that expression levels of voltage-gated potassium channel genes are more likely to show positive than negative relationships with AHP amplitude, whereas the results obtained using the class-independent approach do not appear to support this conclusion.

### Evidence of causal support for specific gene-property relationships

To further validate the gene-property correlations found in the AIBS dataset, we asked whether any of the same relationships showed direct support in the literature. In some cases we found that previously published work showed that manipulation of the gene of interest caused electrophysiological effects in line with what would be predicted by our analysis.

*Kcnal,* a voltage-gated potassium channel, is significantly related to a number of electrophysiological features in our analysis, including maximum firing frequency (FDR= 0.0002; Fig 7A). This finding of a relationship between *Kcna1* expression and maximum firing frequency is consistent with a published study on the same gene. Kopp-Scheinpflug et al. (2003) examined mice with a knockout of the *Kcna1* gene and found that firing rates in auditory neurons were reduced in the knockouts only at high intensities of an auditory stimulus, and that this difference was more robust in the inhibitory neurons of the medial nucleus of the trapezoid body (MNTB) compared to excitatory ventral cochlear nucleus (VCN) bushy cells (7).

**Fig 7.**
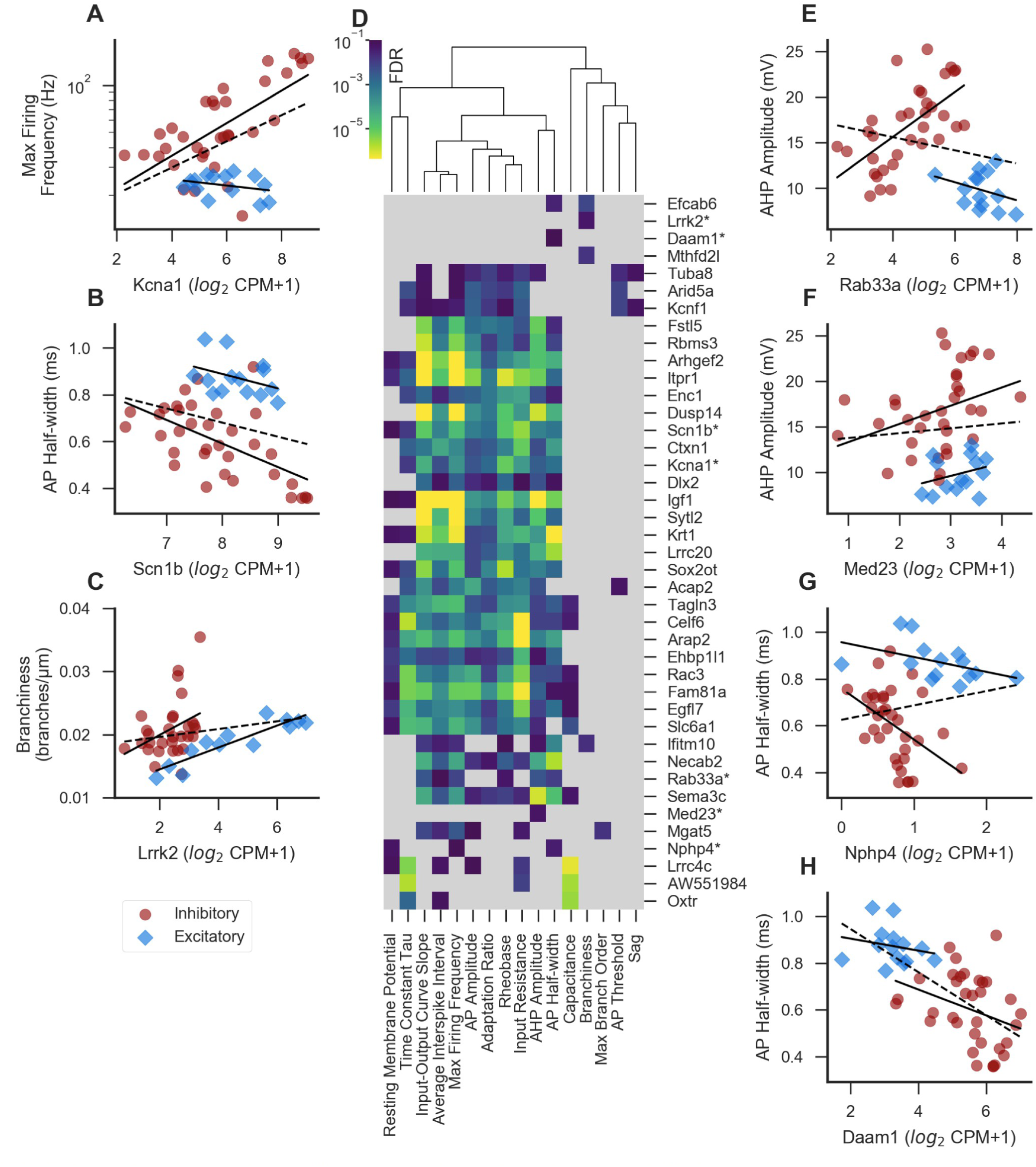
Examples of experimentally supported or otherwise potentially interesting genes. A-C. Examples of genes showing statistica lly-signi ficant gene-property relationships in the class - conditional model (FDR < 0.1) that also have experimental support for their causal regulation of the property in the literature. Solid lines indicate linear fits including only excitatory or only inhibitory cell types, and dashed line indicates a linear fit including all cell types (also applies to E-H). D. Heatmap showing a subset of the most significant genes for each property in the class-conditional model, sorted along both axes by similarity. Dendrogram represents cross-property similarity between the significance levels for the genes shown here; properties appearing closely linked in the dendrogram are those which are strongly associated with the same genes in our analysis. For each property, up to 3 top genes were chosen that were significant (FDR < 0.1) in the class-conditional model, and also non-significant (FDR > 0.2) in both the class-independent and interaction models for the same property. In addition, genes marked by asterisks are shown here based on their known function based on the literature in addition to at least one significant result in the class-conditional model, shown as scatterplots in A-C and E-H. Light grey indicates a non-significant result in the class-conditional model (FDR > 0.1). E-H. Examples of under-studied but plausibly causal genes showing significant results in the class-conditional model (see text).

Expression of *Scn1b*, a voltage-gated sodium channel subunit, shows a negative relationship with action potential half-width in the class-conditional model (FDR = 0.0008; Fig 7B), as well as a number of other properties. This relationship is obscured in the class-independent model due to overall longer half-widths in excitatory cell types. Consistent with the idea that *Scn1b* might function to shorten AP half-widths, layer 5 cortical pyramidal neurons from mice lacking the *Scn1b* gene show longer half-widths than controls, due to changes in protein stability of voltage-gated potassium channels (33).

Interestingly, the *Lrrk2* gene, mutations in which contribute to Parkinson’s disease (34), is positively correlated with neurite branchiness (number of branch points per μm) in the class-conditional model, but not the class-independent model (FDR = 0.046; Fig 7C). *Lrrk2* has been shown by several studies to regulate neurite outgrowth and branching in cultures (35–38).

Not only do the genes discussed here provide important validation for our method, but the existence of a smooth correlation between these genes and their associated properties is potentially interesting. The previous studies cited above provide causal evidence for gene-property relationships via gain- and loss-of-function approaches, which are likely more reminiscent of pathological states than of natural variability between cell types. Our results suggest that these genes could additionally play an instructive role in setting the precise levels of electrophysiological or morphological properties between cell types under normal physiological conditions. In addition, since morphological features are in part established due to developmental gene expression patterns (39), such features may show poor correlations with mRNA sampled from adult cells.

### Novel gene-property relationships

In addition to those discussed above, we identified many genes whose function in regulating neuronal electrophysiology or morphology is less well characterized. These present testable hypotheses for future study. In Table 2, we list some of the top significant genes from the class-conditional model for each property, chosen based on significance levels and/or previous studies into their cellular function (also shown in Fig 7D).

**Table 2.**
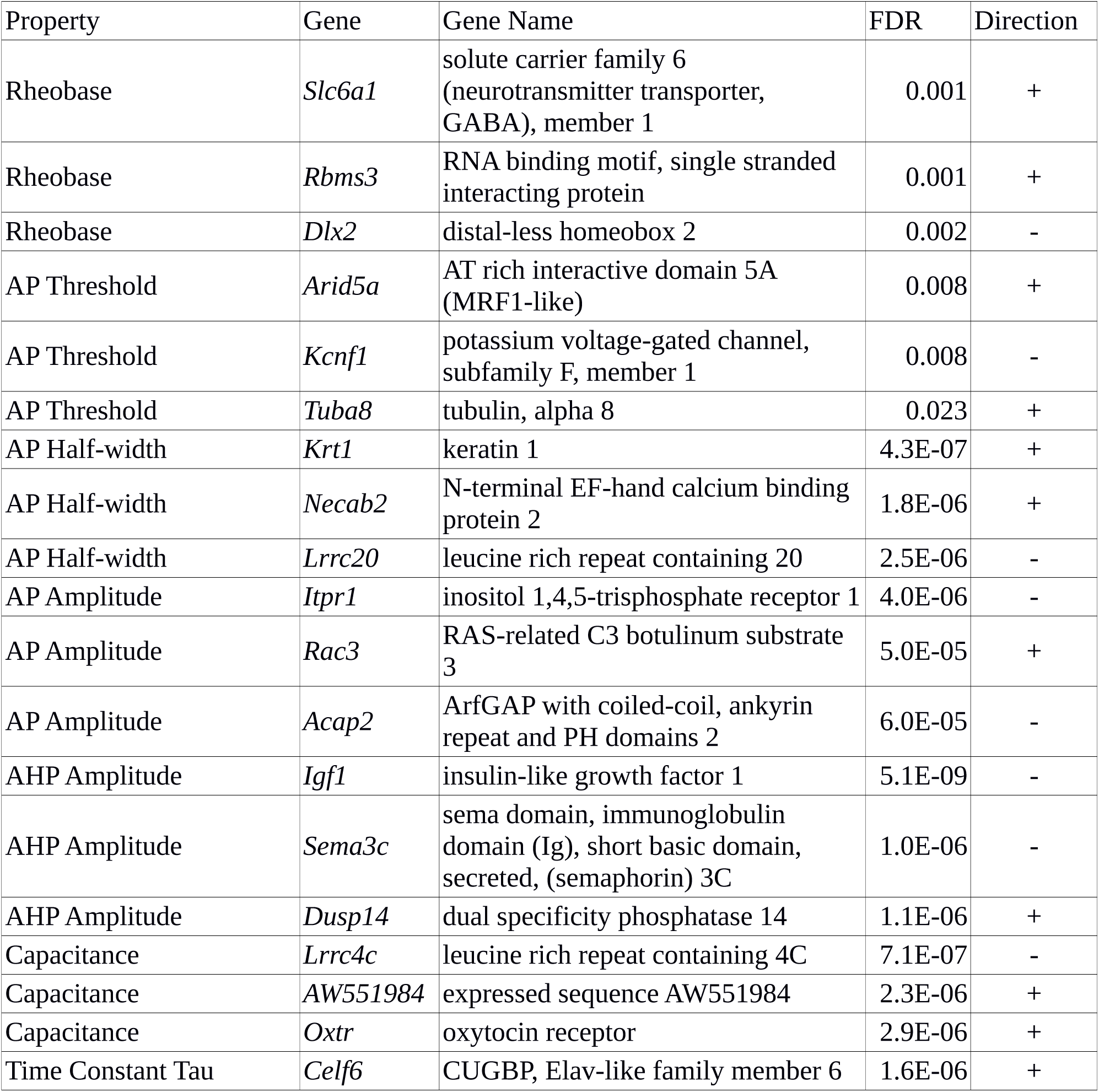
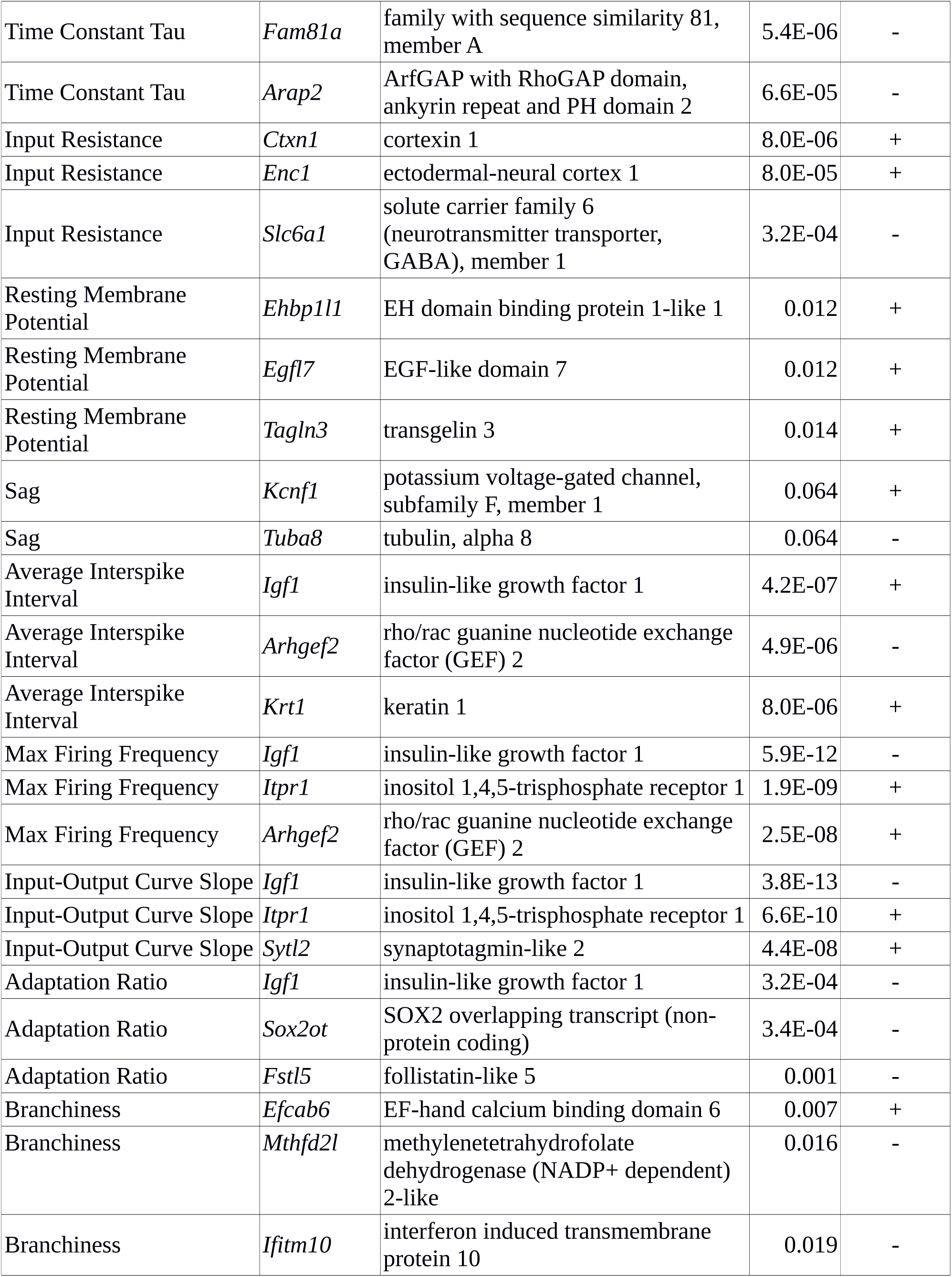
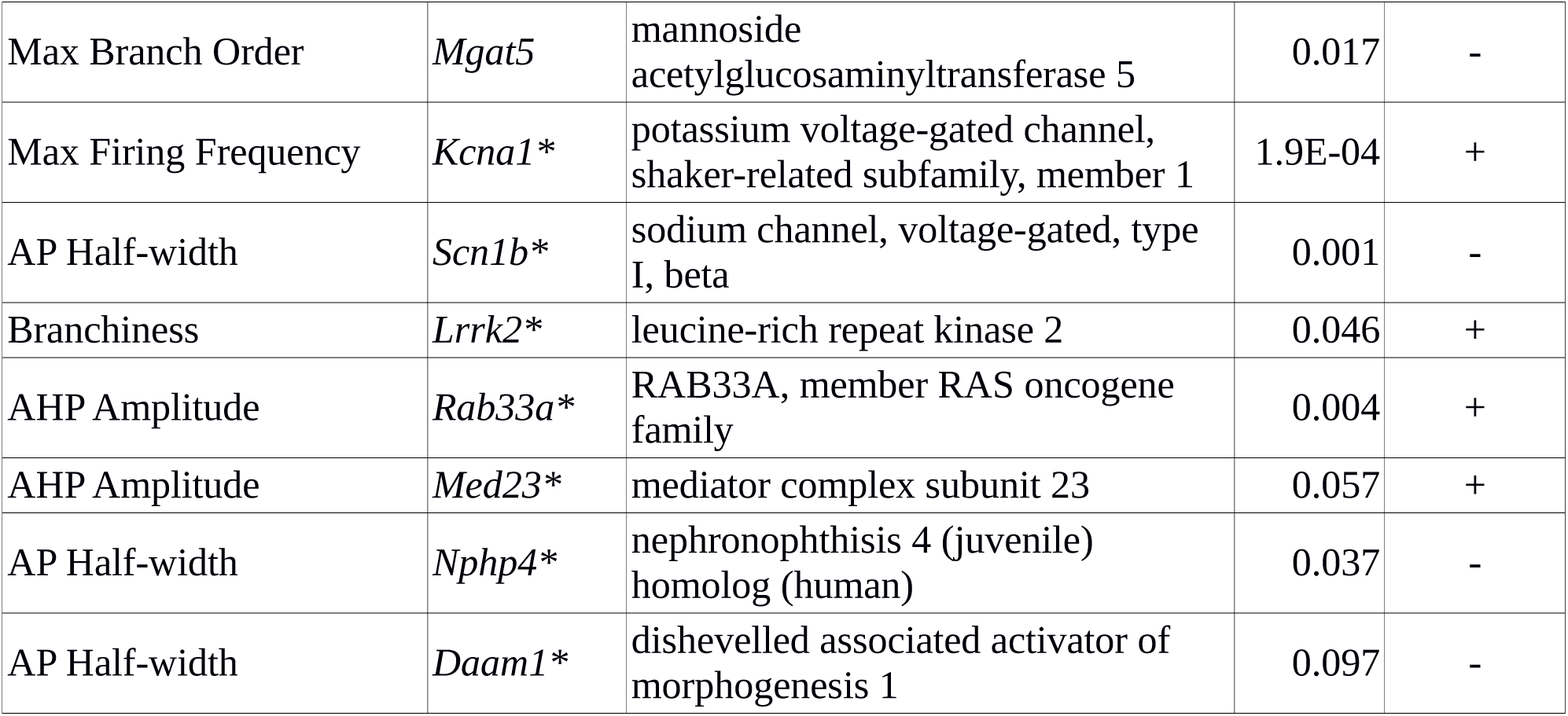
Top correlated genes for each electrophysiological property. Genes marked with asterisks are significantly associated (FDR < 0.1) with the indicated property in the class-conditional model, and selected based on their reported function in the literature. All other genes are significant (FDR < 0.1) in the class-conditional model and non-significant (FDR > 0.2) in both the class-independent and interaction models for the indicated property. “Direction” indicates the direction of the model slope; for example, high expression of Daam1 in a cell type predicts a low value of AP half-width and vice versa.

One notable feature from this analysis is that many of these genes, like *Kcna1* and *Scn1b* discussed above, are significantly associated with several or many different properties. For example, maximum firing frequency, input-output curve slope, and average interspike interval show a similar pattern in the strength of their association with this set of genes. These features all measure similar aspects of neuronal function (broadly speaking, whether a neuron tends to fire rapidly or not), so it would be surprising if they did not show correlations with the same genes. Two more properties that closely share associated genes are AP half-width and AHP amplitude, which measure distinct aspects of the action potential waveform, but might share genetic underpinnings related to rapid channel opening and closing (40). The genes most strongly associated with various electrophysiological properties tend not to show significant associations with the morphological properties of branchiness and max branch order. However, some of the genes associated with these morphological properties do show some (generally weak) associations with some electrophysiological properties (for example *Mgat5* and *Ifitm10*).

Several of the genes for which we were unable to find conclusive loss-of-function studies in the current literature (Fig 7E-H) seem particularly intriguing, given what is known about their cellular function. In the discussion, we briefly speculate about how these genes might function as regulators of the properties with which they are associated in our analysis. However, further study will be needed to determine what role, if any, these genes play in regulating electrophysiological or morphological properties.

## Discussion

In this work we presented a series of correlations between gene expression and electrophysiological or morphological properties, each representing a testable hypothesis for future studies. Our key insight here is to introduce cell class (i.e., excitatory and inhibitory cell type identity) as an indicator variable when modeling the relationship between genes and properties. This has the advantage of 1) avoiding the identification of class-driven correlations, 2) helping identify a subset of non-class-driven correlations that might have been obscured by overall differences between excitatory and inhibitory cell types, and 3) revealing instances where gene-property relationships might be different for excitatory versus inhibitory cell types.

Although the idea that non-class-driven correlations would have a higher chance of being biologically relevant compared to class-driven ones seems straightforward, we evaluated this prediction through a number of specific empirical tests. First, we found better correspondence between gene-property relationships from the class-conditional model with those derived from the non-projecting cell type subset of our prior NeuroExpresso/NeuroElectro dataset. Second, we observed consistency between the class-conditional model and gene-property relationships derived from five independently-collected Patch-seq datasets, suggesting that the relationships described here might be predictive of gene-property relationships within narrowly-defined cell types. Third, our analysis of the relationship between action potential after-hyperpolarization (AHP) amplitude and voltage-gated potassium channel genes suggests that genes and electrophysiological features showing a significant result in the class-conditional model are more likely to reflect known functions of those genes.

The Patch-seq and voltage-gated potassium channel analyses highlighted distinct advantages of the class-conditional model. The class-conditional model revealed higher overlap between the Patch-seq and AIBS datasets, compared to the class-independent model, where most shared relationships (for both models) were in a consistent direction. This indicates that the class-conditional model might be more sensitive to certain relationships, which have some evidence for their biological relevance. In contrast, the main advantage of the class-conditional model in the voltage-gated potassium channel analysis was primarily to avoid class-driven correlations. In other words, the class-conditional model exhibits increased specificity, an important factor when considering that these results might be used to help prioritize genes for experimental study.

In this work, we have operationalized the concepts of class-driven and non-class-driven correlations as those which produce a significant result in the class-independent model only or in the class-conditional model, respectively. This is a simplification, since both effects can exist simultaneously to differing degrees (for example, *Daam1* and AP half-width, Fig 7H) and our ability to distinguish them with confidence is limited by the number and composition of cell types in the dataset. It should be emphasized that, since these categories are defined based on significance thresholds, the distinction between, for example, a non-class-driven relationship which is obscured by class and one which is significant in either model is not meaningful in a statistical sense and should not be interpreted as being directly informative about the underlying biology. Bearing this in mind, the distinction may be useful in practice for prioritizing genes for further examination. Thus, we have shown that thresholding the set of all genes based on one model or the other results in the identification of a distinct but overlapping set of genes, meaning that the choice of model is consequential.

A novel feature of our analysis is the investigation of gene-property relationships that are divergent within excitatory and inhibitory cell types. Using the interaction model, we found a small subset of genes showing significant associations in the class-conditional model that also have a significant interaction term, indicating that their relationship with the property in question is dependent on cell class. We also found another small set of gene-property relationships that have a significant term in the interaction but not the class-conditional model. In contrast to all other properties analyzed, for the properties sag and maximum branch order, the interaction model identified many more genes compared to the class-conditional model. One possible explanation is that for both of these features, the absolute slopes in excitatory cells tend to be higher than those in inhibitory cells (shown in Fig 3B for maximum branch order), suggesting either that these features might be under stronger genetic control in excitatory types compared to inhibitory, or that the genes associated with them in excitatory cell types are more readily identified by our analysis. Since this dataset contains more inhibitory than excitatory types, an inhibitory-specific relationship may be identified in the class-conditional model by virtue of the number of cell types, but an excitatory-specific relationship would likely be “diluted” by the larger number of inhibitory cell types not showing the relationship. It is also possible that, in the case of maximum branch order, this effect is partially explained by methodological differences in the dataset, since inhibitory but not excitatory morphological reconstructions contain axons in addition to dendrites (1).

### Novel putative gene/electrophysiology relationships

Our primary motivation for comparing gene expression to neuronal properties is to identify candidate genes that might influence those properties. While directly testing the functional relevance of specific gene-property predictions is beyond the scope of this work, we have highlighted below some of our potentially novel findings that might be of greatest interest for further follow up.

*Rab33a* expression is positively correlated in the AIBS dataset with AHP amplitude with a significant interaction (Fig 7C), and also shows significant positive correlations with input-output curve slope, maximum firing frequency, and rheobase, and significant negative correlations with AP half-width and average interstimulus interval (ISI). *Rab33a* is a small GTPase thought to be involved in regulation of vesicle trafficking, likely at stages prior to plasma membrane docking (41,42). One hypothesis for how *Rab33a* could regulate AHP amplitude and/or AP half-width is that *Rab33a* might facilitate the transport and/or insertion of vesicles containing voltage-gated potassium channels, or regulators thereof, into the axonal membrane, leading to narrower action potentials and larger AHPs. Our analysis of the AIBS data suggests that any effects of *Rab33a* expression on AHP amplitude would be present only in inhibitory cell types.

*Med23* (also known as *Crsp3*), a subunit of the mediator complex which acts as a transcriptional co-activator for RNA polymerase II (43,44), shows a positive correlation with AHP amplitude (Fig 7D). Although the complete set of roles played by *Med23* are incompletely understood, it has been shown to modulate signaling by the BMP, Ras/ELK1, and RhoA/MAL pathways (45,46). Thus it has the potential to regulate a variety of genes, including potentially voltage-gated potassium channels or interacting proteins thereof. Given *Med23*’s role in regulating transcription through a variety of signaling pathways, it is notable that our analysis showed only one feature with which it was convincingly associated. It is also interesting to note that mutations in *Med23* have previously been associated with intellectual disability, in some cases with a predisposition to seizures (47,48).

Expression of *Nphp4* encoding the cytoskeletal-associated protein nephrocystin-4 was negatively correlated with AP half-width (Fig 7E) as well as with resting membrane potential and maximum firing frequency. Although *Nphp4* is primarily understood for its function in the kidney, *Nphp4* mutations often cause co-morbid deficits in the nervous system (49). Furthermore, *Nphp4* has been shown to regulate actin networks via its interaction with the polarity protein *Inturned* and with the formin *Daam1* (50). *Daam1* is also negatively correlated with AP half-width (Fig 7F), and not significantly correlated with any other features. The actin network in the axon forms a highly regular lattice structure which includes regularly interspersed voltage-gated sodium channels (51). A similar relationship between the actin network and other voltage-gated ion channels has not been tested, but seems plausible. A potential mechanism through which *Nphp4* and *Daam1* could regulate the shape of the action potential might involve the organization of the axonal actin network structure, which might change the local levels or relative positioning of voltage-gated ion channels, especially potassium channels, or their regulators.

### Limitations and Caveats

We note that the gene-property relationships reported here are by definition correlational. Demonstrating that any specific gene is involved in regulation of any electrophysiological or morphological property is beyond the scope of this work. Our goal in this study was to generate testable hypotheses which, together with the current body of published literature, will help guide future experiments. We expect that this list of putative relationships contains some proportion of causal genes, and based on our analyses expect that this proportion may be higher than that in our previous work (11), However, causality can only be determined for a given gene and property using direct experimental methods.

Additionally, as in our prior work (11), we have limited our analyses to models in which expression levels of a single gene predict downstream properties in an approximately linear fashion, and in which that gene is regulated primarily at the transcriptional level. Some instances of mechanisms involving interactions between multiple genes, or those involving a non-linear relationship between log-gene expression and an electrophysiological or morphological property, are likely to have been missed here. In addition, for mechanisms through which electrophysiological or morphological properties are controlled at the translational or post-translational level, our analysis is unlikely to provide insight into the gene whose product directly controls the property. However, this analysis has the power to identify transcripts whose products are involved in the translation, modification, or trafficking of proteins which in turn regulate electrophysiology or morphology.

Furthermore, the generalizability of the gene-property relationships reported here might be limited by the fact that the AIBS dataset only reflects cells sampled from the adult mouse primary visual cortex. Therefore, the relevance of our results to other brain regions depends on the assumption that many of the same genes regulate electrophysiological or morphological properties in different cell types. This assumption of generalizability across brain areas appears to be appropriate in the case of *Kcna1* and maximum firing frequency (Fig 7A and (7)). Additionally, this assumption is supported by our comparisons with the NeuroExpresso/NeuroElectro dataset and Patch-seq datasets, both of which contain cells sampled from other brain regions. However, some relationships may not generalize across brain regions due to differences in expression of other genes or the presence of post-translational modifications which modify the consequences of expressing a given gene.

Another potential confounding factor in our reliance on the AIBS datasets is the uneven balance in the count of inhibitory versus excitatory cell types. The practical consequence of this is that the results from the class-conditional model are likely biased towards explaining gene-property relationships within inhibitory cell types, and might be missing relationships that are specific to excitatory cell types. Even in the absence of a significant interaction term, gene-property relationships may have stronger evidence in one cell class than the other. An example of this is *Lrrk2* and branchiness (Fig 7C), where despite very similar slopes between classes and no statistical evidence of an interaction, the correlation among excitatory cells is much tighter than that among inhibitory cells. For this reason, when prioritizing genes for future study, we strongly recommend making a plot of gene, property, and cell class before concluding that the overall result is likely to apply to both classes.

### Future Directions

The primary goal of this project was to produce a list of genes which we can recommend for future study based on their correlations with electrophysiological and morphological properties in the AIBS dataset. We believe that some of the genes we identified are promising candidates for future study.

In order to facilitate the use of our results by others in prioritizing genes for investigation, we are providing a Jupyter Notebook file to facilitate exploration of the data (available at https://github.com/PavlidisLab/transcriptomic_correlates). We have endeavored to make this easy to use for researchers with little or no coding experience. We encourage those who are interested in a particular electrophysiological or morphological property, gene, or set of genes, to explore the data and to make their own judgements as to which genes are worth following through on experimentally and which measures should be prioritized for recording. Our recommendation is to use the gene list in conjunction with other sources of information about gene function, such as Gene Ontology annotations (52,53) and previously published literature, in prioritizing genes for future study.

## Materials and Methods

### AIBS Dataset

The RNA-seq dataset from (14) was accessed via the Allen Institute for Brain Science’s Cell Types database (http://celltypes.brain-map.org/) on June 19, 2018, and contains 15,413 cells isolated by microdissection and fluorescence-activated cell sorting from primary visual cortex of mice expressing tdTomato under the control of various Cre driver lines. Electrophysiological and morphological data were also accessed via the Allen Institute for Brain Science Cell Types database on June 21, 2018. The dataset includes electrophysiological recordings from 1920 cells, of which 1815 are reporter-positive, from the visual cortex of mice also expressing tdTomato driven by Cre, many of which are from the same lines represented in the RNA-seq dataset. A subset of these cells (509, of which 471 are reporter-positive) have morphological reconstruction data available. Cells in both the electrophysiology/morphology and RNA-seq datasets are annotated according to the cortical layer they reside in (for electrophysiology/morphology this is always a single layer, and for RNA-seq may be a single layer, subset of layers, or all layers), their Cre-line, and whether they express the reporter.

### Filtering and matching datasets

Single-cell RNA-sequencing data, summarized as counts per million reads sequenced (CPM), were log2-transformed prior to combining with electrophysiological and morphological data. Cells from the RNA-seq dataset were excluded if they were annotated as having failed quality control checks, if they were negative for expression of tdTomato, or if they were labeled as non-neuronal or unclassified. Cells in the electrophysiology/morphology dataset were excluded if they were negative for expression of tdTomato.

### Electrophysiological and morphological measures

Electrophysiological data were downloaded from http://celltypes.brain-map.org/ and summarized as described previously (11) except for the features response frequency versus stimulus intensity (input-output) curve slope, average interstimulus interval (ISI), and sag, which we did not use previously as they were not represented in the NE dataset. All three of these new features were pre-computed in the downloaded dataset. In order to include only sag values which could be meaningfully compared, any cells having a value of “vm-for-sag” (the membrane voltage at which sag values were measured) not between −90 and −110 mV, or having a resting membrane potential lower than −80 mV, were excluded from analyses of sag, but were used for analyses of other electrophysiological features. The morphological features “average_bifurcation_angle_local”, “max_branch_order”, “soma_surface”, “total_length”, and “total_volume” were pre-computed in the dataset. We defined “branchiness” according to the pre-computed feature “number_branches” divided by “total_length” as a measure of how often a given cell produces branches per unit of neurite length. For the features input resistance, tau, capacitance, rheobase, maximum firing frequency, AHP amplitude, adaptation ratio, input-output curve slope, latency, branchiness, max branch order, total length, and total volume, values were log10-transformed prior to use in order to mitigate underlying skew or non-normality in these data values.

### Defining cell types

Cell types in the AIBS dataset were defined according to the Cre-line they were isolated from, whether they were excitatory or inhibitory, and in most cases either a single cortical layer or a range of layers. Where multiple layer dissections containing a sufficient number of cells were present for a Cre-line in the RNAseq data, we decided on whether and how to combine layers based on the following criteria: 1) producing the maximum number of cell types, 2) producing the most homogenous cell types possible, and 3) producing cell types containing sufficiently large numbers of cells in both the RNA-seq and electrophysiology or morphology datasets. The first two criteria favored splitting layers more finely, whereas the last favored combining layers. Only cell types where both datasets contained at least 6 cells (for the electrophysiology analysis) or at least 3 cells (for the morphology analysis) were included in the final analysis. Cell type definitions, along with the numbers of cells meeting the criteria for each type, are shown in table S1.

Splitting cells from certain Cre-lines into multiple types based on their layer location and their identity as excitatory or inhibitory allowed us to increase the number of types in our analysis. Splitting cell types in this way makes biological sense in that cells isolated from the same Cre-line but different layers often belong to different transcriptomically-defined cell types. For example, cells isolated from from the upper cortical layers of Sst-Cre mice primarily belong to the Sst Cbln4 type, whereas the majority of cells from lower layers belong to either the Sst Myh8 or Sst Th types (15). We have further justified this decision based on the fact that there are frequently electrophysiological differences between cells from the same Cre-line but from different layers (examples of three electrophysiological properties are shown in Fig S1).

After the two datasets were matched, the combined dataset contained 1359 cells belonging to 48 types with electrophysiological data, 369 cells belonging to 43 types with morphological data, and 4403 cells belonging to 50 types with RNA-seq data (Table S1). The remaining cells in the original datasets were those whose types could not be matched, either because the Cre-line or layer they were isolated from was not sampled in the other datasets, or because the number of cells belonging to that type was below our threshold for the number of cells per type required.

### Modeling the relationship between gene expression and electrophysiology/morphology

Mean expression values for each gene and mean values for each electrophysiological or morphological property were calculated for each cell type as defined above. If more than two cell types showed zero expression of any given gene, those cell types were removed from analyses for that gene. We found this step to be necessary in initial analyses because differences in electrophysiology/morphology among these cell types could not be assessed in relation to differences in gene expression, potentially producing spurious correlations. Any genes for which this left fewer than eight samples were excluded. Out of all genes represented in the RNA-seq dataset, ~26% passed this thresholding step. For the remaining genes, and for each electrophysiological or morphological property, we fit one or more linear models relating the property (P) to expression of the gene (G) and/or cell class (C). Model 1 (P~G; “class-independent model”) attempted to explain the property based on only expression of the gene. For genes which were expressed in both excitatory and inhibitory types, we fit three additional models. Model 2 (P~C) related property to cell class, model 3 related the electrophysiological parameter to the gene and cell class (P~G+C), and model 4 related the electrophysiological parameter to gene, cell class, and an interaction term between gene and cell class (P~G+C+G*C). Models 2 and 3, as well as models 3 and 4, were compared to one another using an ANOVA, resulting in the “class-conditional model” (P~G|C) and “interaction model” (P~G*C|G+C), respectively. Beta coefficients from models 1, 3, and 4 (separately for each cell type) were recorded, as well as p-values from model 1 and from both ANOVAs. Prior to filtering for significantly-correlated genes, false discovery rate (FDR) correction was performed using the Python package statsmodels.sandbox.stats.multicomp.fdrcorrection0 with an alpha level of 0.05. Model 2 was also used directly to test for significant differences between cell classes in the value of each property.

### Non-projecting class-specific correlations in the NeuroElectro/NeuroExpresso dataset

The NeuroElectro and NeuroExpresso datasets were described previously (11). In order to limit the dataset to only non-projecting cell types (13), we chose cells whose major type was annotated as anything other than “Pyramidal,” “Glutamatergic,” or “MSN”. Cells of the types “Ctx Htr3a” and “Ctx Oxtr” were excluded due to their lower transcriptomic quality compared to others in the dataset (54). After subsetting, 19 cell types remained. Average values were calculated for gene expression and electrophysiological properties across cells within a type, and Spearman correlations were calculated for each combination of gene and electrophysiological property.

In order to assess cross-dataset consistency, we calculated a Spearman correlation between the beta coefficients (slopes) resulting from the class-independent or class-conditional model in the AIBS dataset and the correlation values calculated in the NE dataset. If there was a significant positive correlation between the AIBS slope and the NE correlation value, we concluded that the results of the two analyses were consistent (although this does not imply that they were highly consistent). For those comparisons which were consistent, we considered one method to be “more consistent” than the other if the AIBS/NE correlation value was higher (with non-overlapping 95% confidence intervals) than that derived using the second method.

### Data Analysis and Visualization

All statistical analyses and data visualization were performed using Jupyter Notebook (55) and Python 2.7, and the following packages: scipy.stats, numpy, pandas, matplotlib, mpl_toolkits, matplotlib_venn, seaborn, statsmodels.sandbox.stats.multicomp.fdrcorrection0, mygene.

Bootstrapped confidence intervals and significance between models for correlations between the NE and AIBS datasets were calculated as follows: Starting with the list of paired correlation values and beta coefficients for a given electrophysiological feature and model (class-independent or class-conditional), in which each pair represented a single gene and each value in that pair was calculated using one of the two datasets, a new list of paired correlation values of the same length was calculated by resampling with replacement. A new Spearman correlation was then calculated based on the resampled list. The resampling procedure was repeated 100 times, and the upper and lower ends of the confidence intervals were calculated by finding the values at the 2.5th and 97.5th percentiles. Significance was determined by finding the difference between each pair of resampled correlations from the two models, and then again finding the values at the 2.5th and 97.5th percentiles. If this interval did not contain zero, the two consistency metrics were said to be significant at p < 0.05.

Hierarchical clustering in Fig 7D was performed using the seaborn.clustermap tool using the “average” (UPGMA) method and the euclidean metric (56,57).

### Data Availability

Analysis code and processed data will be available at https://github.com/PavlidisLab/transcriptomic_correlates. Included there is a Jupyter notebook file with some recommended steps for filtering and visualizing results, which can be run directly from the user’s web browser without any need for installation of software. We have made an effort to make this resource approachable for researchers with little or no coding experience. The Bengtsson Gonzales Patch-seq dataset will be made publicly available.

### Analysis of Patch-seq datasets

#### Overview of datasets used

Our analysis of the Patch-seq datasets builds on our analysis described previously (21). Here, we made use of four previously published Patch-seq datasets that have characterized interneurons of the mouse forebrain, described in detail in Table 1. (“Cadwell,” “Földy,” “Fuzik,” “Muñoz”; (17,22,26,27)). Our analysis also includes one novel dataset of 19 Pvalb-Cre positive interneurons recorded in region CA1 of the mouse hippocampus, reported here for the first time. Cells in this dataset (referred to as the Bengtsson Gonzales dataset), were treated, processed, and analyzed using the same methodology as described in (22).

Datasets were processed and normalized as described in (21) with a small number of exceptions. First, datasets employing unique molecule identifiers (UMIs), including the Fuzik, Muñoz and Bengtsson Gonzales datasets, were normalized to a total library size of two thousand UMIs per cell. Similarly, the Cadwell and Földy datasets were normalized to counts per million (CPM), to be more directly comparable with how we have normalized the AIBS datasets here. Second, because Patch-seq sampled cells varied considerably in amount of mitochondrial and other non-coding mRNAs, when normalizing cells to the total count of reads detected in each cell, we only quantified reads mapping to protein coding genes, as defined by biomaRt (58). Furthermore, we used biomaRt to help reconcile gene names between Patch-seq datasets.

#### Assigning Patch-seq single cells to transcriptomically-defined cell types

We implemented a nearest-centroid classifier to map Patch-seq transcriptomes to transcriptomically defined clusters, as defined in the Tasic 2018 cortical and Muñoz-Manchado 2018 striatum reference atlases. Specifically, for each transcriptomically-defined cluster in these reference datasets, we first calculated the mean expression level across all cells assigned to the cluster. Next, using the two thousand most variable genes amongst inhibitory cell types in the Tasic dataset (described in the section below), we calculated the Spearman correlation of each Patch-seq cell to every cluster in the dissociated cell dataset and assigned cells to the cluster that they were most correlated with (we compared all Patch-seq datasets except the striatum Muñoz dataset to the Tasic cortical dataset). For cortical and hippocampal cell types, to increase the number of cells defined per transcriptomic type, we made use of the ‘subclass’ mappings provided in the Tasic 2018 dataset, mapping neurons to the Pvalb, Sst, Vip, Lamp5, and Sncg major interneuron cell types. To estimate transcriptome quality we used the “quality score” metric from our prior analysis, using the full set of “on” and “off” marker genes.

#### Identifying highly variable genes per cell type

We used the ‘decomposeVar’ function from the ‘scran’ R package (59) to identify highly variable genes in each subclass in the Tasic 2018 dataset and each cell type in the Muñoz-Manchado reference datasets.

#### Mixed effects statistical model to identify gene-property relationships in Patch-seq cell types

We used a mixed effects model of the following form with gene expression as a fixed effect and dataset and cell type as random effects:

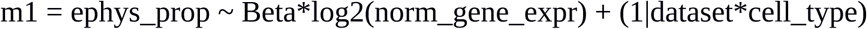

where we used an anova to test for the significance of the beta associated with the gene expression term by comparison to an equivalent statistical model without the gene expression term. We used the quality score as a weight in the regression analysis, and normalized these across datasets. We used the ‘lmer’ function within the ‘lme4’ R package for fitting mixed-effects models. We performed this analysis on the top 250-most variable genes per cell type and for genes that were highly variable in at least one cell type across at least 2 (of the 5 total) Patch-seq datasets used here. In addition, we did not use Patch-seq cell types where gene expression was detected in fewer than 33% of cells and with fewer than 5 cells expressing the gene.

## Acknowledgements

Some of the ideas explored in this work resulted from the Allen Institute for Brain Science 2017 Summer Workshop on the Dynamic Brain. We would like to thank the organizers, instructors, and participants in this event for their guidance and support. In particular, we would like to thank Agata Budzillo and Nathan Gouwens for their helpful advice and guidance, and Annie Vogel-Cierna for her invaluable input at the early stages of the project, as well as for providing feedback on the manuscript. We would also like to acknowledge the investigators responsible for collecting the data represented in the AIBS Cell Types Database as well as the Patch-seq datasets re-analyzed in this study. We would like to thank the Eukaryotic Single Cell Genomics facility at Science for Life Laboratory.

PP was funded by Kids Brain Health Network, Natural Sciences and Engineering Research Council Discovery grant RGPIN-2016-05991, and NIH grant MH111099. SJT was funded by a Canadian Institute for Health Research Post-doctoral Fellowship. AMC was supported by CIHR FDN-143206 and Canada Research Chair. JH-L was funded by the Swedish Research Council (Vetenskapsrådet, award 2014-3863), StratNeuro, and the Swedish Brain Foundation (Hjärnfonden). CBG was funded by the NIH-KI doctoral program. The funders had no role in study design, data collection and analysis, decision to publish, or preparation of the manuscript.

## Author Contributions

CB, SJT, and PP conceived the project. CB and SJT performed the AIBS and Patch-seq analyses, respectively. CBG collected some of the data used in the Patch-seq analysis (Bengtsson Gonzales dataset) under the supervision of JH-L. CB and SJT wrote the original draft of the manuscript, and all authors contributed to review and editing.

## Competing Interests

The authors declare no competing financial interests.

## Supporting Information

**Fig S1.**
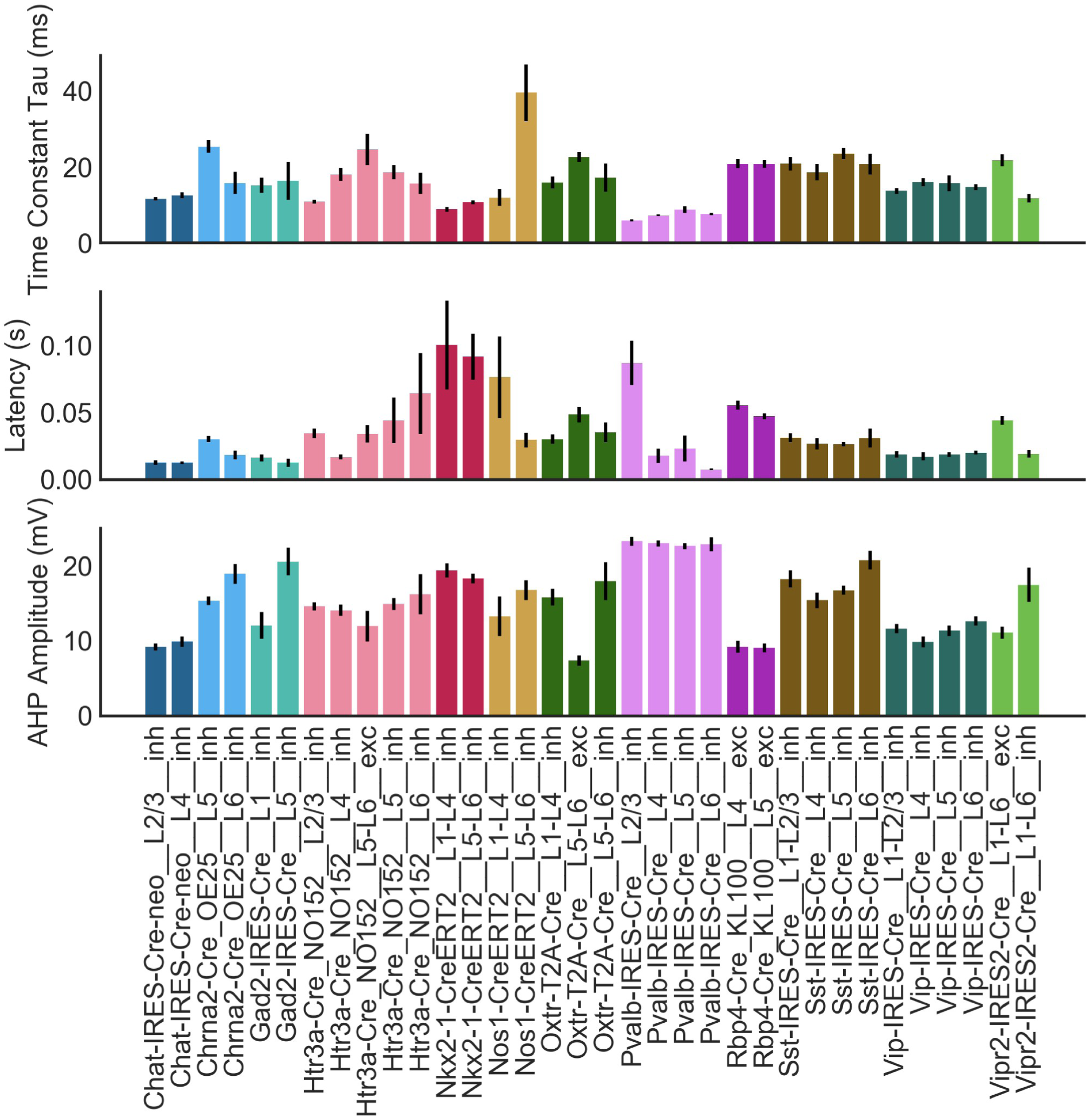
Justification for cell type definitions in the AIBS dataset. Cell types defined based on the same Cre line but different layers and/or excitatory/inhibitory identity show differences in eletrophysiological features. Data are represented as mean ± SEM.

**Fig S2.**
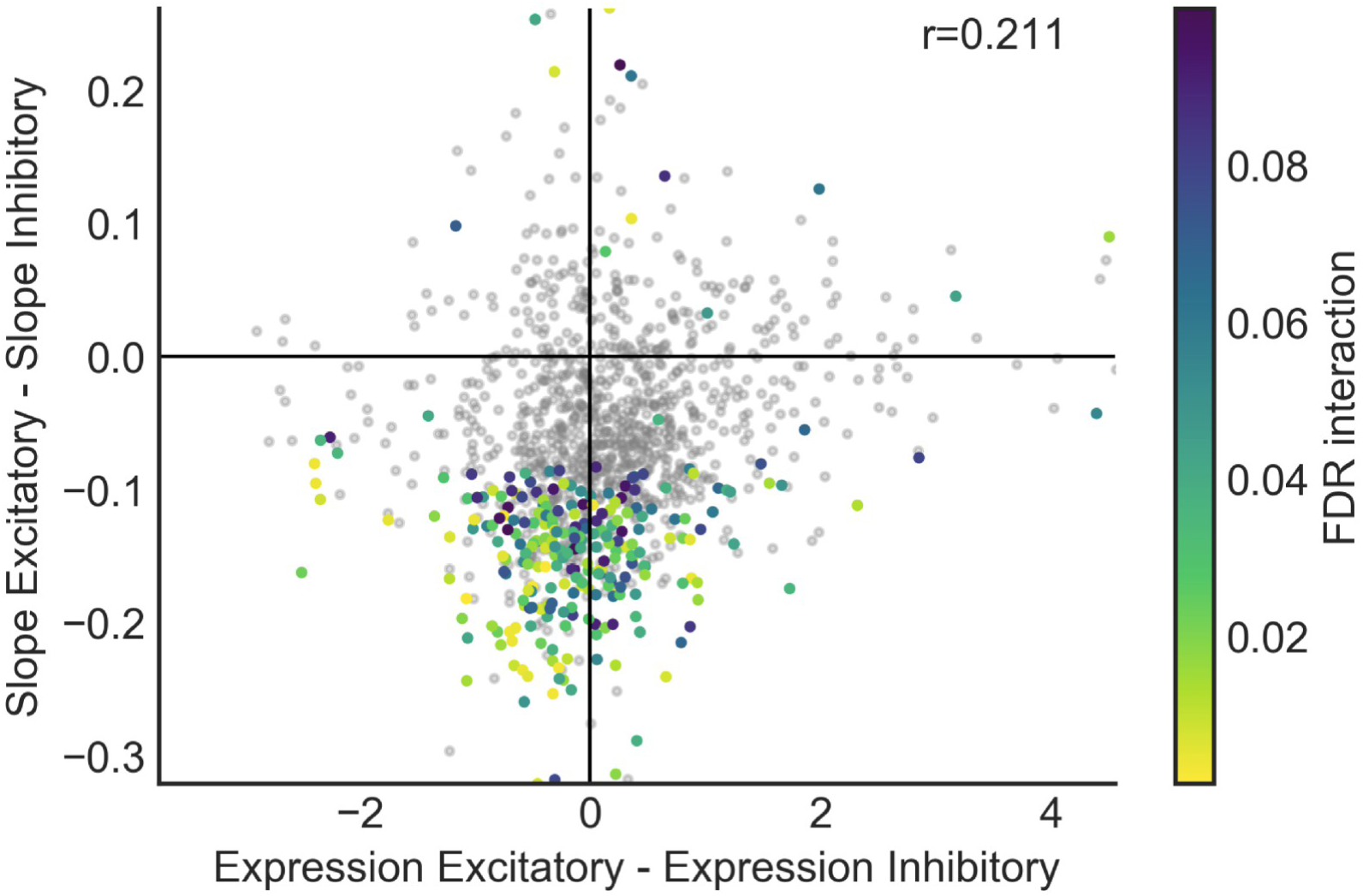
Interactions do not result primarily from low gene expression in one cell class. Between-class differences in gene expression plotted against differences in gene-property slope in the interaction model for the property AHP amplitude. Each point represents a single gene; grey points do not have a significant interaction and others are colored according to their significance level in the interaction model. For clarity of visualization only a random subset of the data (10% of the total number of genes) are plotted.

**Table S1.** Criteria used for defining cell types from the AIBS dataset according to the cre line and layer they were isolated from as well as excitatory/inhibitory identity. For each cell type, the number of cells meeting the criteria which were profiled for each of the three data modalities are indicated. For electrophysiology and morphology, blank cells indicate that not enough cells meeting the criteria were present in that dataset, so that cell type was not included in the analysis.

|  |  |  | RNA-seq | Electrophysiology | Morphology |
| --- | --- | --- | --- | --- | --- |
| Chat-IRES-Cre-neo | L2/3 | inh | 36 | 45 | 15 |
| Chat-IRES-Cre-neo | L4 | inh | 8 | 18 | 6 |
| Chrna2-Cre_OE25 | L5 | inh | 57 | 29 | 9 |
| Chrna2-Cre_OE25 | L6 | inh | 40 | 10 |  |
| Chrna2-Cre_OE25\|Pvalb-T2A-Dre | L5-L6 | inh | 59 | 8 |  |
| Ctgf-T2A-dgCre | L6 | exc | 150 | 43 | 16 |
| Esr2-IRES2-Cre | L5-L6 | exc | 37 | 20 | 3 |
| Gad2-IRES-Cre | L1 | inh | 298 | 7 | 3 |
| Gad2-IRES-Cre | L5 | inh | 327 | 7 | 3 |
| Htr3a-Cre_NO152 | L2/3 | inh | 32 | 70 | 16 |
| Htr3a-Cre_NO152 | L4 | inh | 31 | 21 | 4 |
| Htr3a-Cre_NO152 | L5 | inh | 30 | 39 | 12 |
| Htr3a-Cre_NO152 | L5-L6 | exc | 9 | 7 |  |
| Htr3a-Cre_NO152 | L6 | inh | 23 | 8 | 3 |
| Htr3a-Cre_NO152\|Pvalb-T2A-Dre | L5-L6 | inh | 7 | 10 | 3 |
| Ndnf-IRES2-dgCre | L1 | inh | 52 | 39 | 9 |
| Nkx2-1-CreERT2 | L1-L4 | inh | 32 | 22 | 6 |
| Nkx2-1-CreERT2 | L5-L6 | inh | 13 | 26 |  |
| Nos1-CreERT2 | L1-L4 | inh | 92 | 8 | 4 |
| Nos1-CreERT2 | L5-L6 | inh | 34 | 10 | 3 |
| Nos1-CreERT2\|Sst-IRES-FlpO | L1-L4 | inh | 3 |  | 3 |
| Nos1-CreERT2\|Sst-IRES-FlpO | L5-L6 | inh | 20 | 35 | 13 |
| Nr5a1-Cre | L4 | exc | 243 | 55 | 15 |
| Ntsr1-Cre_GN220 | L6 | exc | 97 | 51 | 16 |
| Oxtr-T2A-Cre | L1-L4 | exc | 22 |  | 4 |
| Oxtr-T2A-Cre | L1-L4 | inh | 117 | 21 | 8 |
| Oxtr-T2A-Cre | L5-L6 | exc | 71 | 14 | 4 |
| Oxtr-T2A-Cre | L5-L6 | inh | 62 | 6 |  |
| Penk-IRES2-Cre-neo | L5-L6 | exc | 44 | 8 | 3 |
| Pvalb-IRES-Cre | L2/3 | inh | 213 | 42 | 13 |
| Pvalb-IRES-Cre | L4 | inh | 53 | 69 | 9 |
| Pvalb-IRES-Cre | L5 | inh | 218 | 83 | 28 |
| Pvalb-IRES-Cre | L6 | inh | 47 | 22 | 10 |
| Rbp4-Cre_KL100 | L4 | exc | 16 | 21 | 4 |
| Rbp4-Cre_KL100 | L5 | exc | 695 | 55 | 20 |
| Scnn1a-Tg2-Cre | L4 | exc | 22 | 32 | 8 |
| Scnn1a-Tg3-Cre | L2/3-L4 | exc | 107 | 58 | 13 |
| Sim1-Cre_KJ18 | L4-L6 | exc | 74 | 32 | 6 |
| Slc32a1-T2A-FlpO\|Vipr2-IRES2-Cre | L1-L4 | inh | 17 | 25 | 10 |
| Sst-IRES-Cre | L1-L2/3 | inh | 80 | 15 | 8 |
| Sst-IRES-Cre | L4 | inh | 11 | 23 | 3 |
| Sst-IRES-Cre | L5 | inh | 140 | 66 | 21 |
| Sst-IRES-Cre | L6 | inh | 125 | 17 | 6 |
| Tlx3-Cre_PL56 | L4-L6 | exc | 115 | 41 | 11 |
| Vip-IRES-Cre | L1-L2/3 | inh | 149 | 32 | 6 |
| Vip-IRES-Cre | L4 | inh | 67 | 16 | 4 |
| Vip-IRES-Cre | L5 | inh | 91 | 17 |  |
| Vip-IRES-Cre | L6 | inh | 38 | 29 | 3 |
| Vipr2-IRES2-Cre | L1-L6 | exc | 43 | 16 |  |
| Vipr2-IRES2-Cre | L1-L6 | inh | 36 | 11 | 5 |
| Number of Types |  |  | 50 | 48 | 43 |

**Table S2.** Overlap between class-independent and class-conditional models. Comparison of the number of genes showing a significant result (FDR < 0.1) for each electrophysiological or morphological property in the class-independent or class-conditional model, and extent of overlap between these two sets of genes. Definitions of electrophysiological properties are reproduced from (11), except for input-output curve slope, latency, ISI CoV, average ISI, and sag, which are described based on the Allen Cell Types database (http://celltypes.brain-map.org/). Morphological features are described based on (1).

|  | Class-independent model | Class-conditional model | Significant in both models | Definition | Units | Transform |
| --- | --- | --- | --- | --- | --- | --- |
| Soma Surface | 0 | 0 | 0 | Surface area of the cell body | $\mu\text{m}^2$ | linear |
| Total Volume | 1019 | 0 | 0 | Volume of the cell, including cell body as well as processes | $\mu\text{m}^3$ | log10 |
| Total Length | 3398 | 0 | 0 | Total length of all processes | $\mu\text{m}$ | log10 |
| Max Branch Order | 4308 | 4 | 3 | Maximum number of times that a process bifurcates between the soma and branch tip |  | log10 |
| Branchiness | 35 | 132 | 35 | Number of bifurcations encountered per process length |  | log10 |
| Bifurcation Angle | 0 | 0 | 0 | Mean angle across all bifurcation points | degrees | linear |
| Adaptation Ratio | 4164 | 3220 | 2220 | Ratio of durations between early and late AP inter-spike intervals in an AP train | ratio | log10 |
| Input-Output Curve Slope | 6424 | 7022 | 4583 | Slope of the relationship between current injection and resulting firing frequency, based on multiple long current steps | Hz/pA | log10 |
| Max Firing Frequency | 6113 | 6320 | 3977 | Maximum observed AP discharge rate | Hz | log10 |
| Latency | 566 | 0 | 0 | Latency to fire the first action potential during a long current step | s | log10 |
| Interspike Interval Coefficient of Variation (ISI CoV) | 30 | 0 | 0 | Variability between interspike intervals within one sweep, measured as standard deviation/mean | ratio | log10 |
| Average Interspike Interval | 5405 | 4447 | 2699 | Average time elapsed between spikes during a sweep | ms | log10 |
| Sag | 0 | 2 | 0 | Measure of the extent to which the membrane potential recovers toward resting potential when the neuron is strongly hyperpolarized (between -90 and -110 mV) | ratio | log10 |
| Resting Membrane Potential | 1546 | 443 | 280 | Membrane potential at the onset of whole-cell recording | mV | linear |
| Input Resistance | 2615 | 3373 | 2404 | Input resistance measured at steady-state voltage response to current injection | $\text{M}\Omega$ | log10 |
| Time Constant Tau | 3204 | 1441 | 1078 | Time constant for the membrane to repolarize after a small current injection of fixed amplitude and duration | ms | log10 |
| Capacitance | 5508 | 1736 | 1144 | Neuron capacitance, typically measured by dividing membrane time constant by membrane resistance | pF | log10 |
| After-hyperpolarization (AHP) Amplitude | 6056 | 6568 | 3681 | Calculated as the voltage difference between AP threshold and AP trough. Commonly defined using first AP in train at rheobase current. | mV | log10 |
| Action Potential (AP) Amplitude | 4969 | 3438 | 1997 | Voltage indicating height of action potential. Usually calculated as the difference between AP peak and AP threshold voltages. Commonly measured using first AP in train at rheobase current. | mV | linear |
| AP Half-width | 5384 | 4522 | 2489 | Calculated as the AP duration at the membrane voltage halfway between AP threshold and AP peak. Most commonly calculated using first AP in train at rheobase current. | ms | linear |
| AP Threshold | 31 | 24 | 11 | Voltage at which AP is initiated (as assessed by measuring rising slope of membrane voltage) | mV | linear |
| Rheobase | 3238 | 4108 | 3237 | Minimum current injected somatically required to fire AP | pA | log10 |

**Table S3.** Overlap between class-conditional and interaction models. Comparison of the number of genes showing a significant result (FDR < 0.1) for each electrophysiological or morphological property in the class-conditional or interaction model, and extent of overlap between these two sets of genes.

|  | Class-conditional model | Interaction model | Significant in both models |
| --- | --- | --- | --- |
| Soma Surface | 0 | 0 | 0 |
| Total Volume | 0 | 0 | 0 |
| Total Length | 0 | 0 | 0 |
| Max Branch Order | 4 | 1914 | 0 |
| Branchiness | 132 | 5 | 0 |
| Bifurcation Angle | 0 | 0 | 0 |
| Adaptation Ratio | 3220 | 325 | 253 |
| Input-Output Curve Slope | 7022 | 408 | 388 |
| Max Firing Frequency | 6320 | 335 | 312 |
| Latency | 0 | 0 | 0 |
| ISI CoV | 0 | 0 | 0 |
| Average Interspike Interval | 4447 | 54 | 47 |
| Sag | 2 | 1174 | 0 |
| Resting Membrane Potential | 443 | 10 | 1 |
| Input Resistance | 3373 | 99 | 89 |
| Time Constant Tau | 1441 | 123 | 103 |
| Capacitance | 1736 | 156 | 101 |
| AHP Amplitude | 6568 | 2962 | 2222 |
| AP Amplitude | 3438 | 658 | 457 |
| AP Half-width | 4522 | 96 | 73 |
| AP Threshold | 24 | 4 | 0 |
| Rheobase | 4108 | 91 | 77 |

The following are in separate files:

*Table S4. Table of all significant results*

*Correlation and significance values for all combinations of gene and electrophysiological and morphological features which were significant at FDR <0.1 in either the class-conditional, the interaction model, or both. Each entry is annotated with the total number of features for which the same gene was significant at padj <0.1 as a measure of the extent to which that gene is either unique to that feature or shared between features.*

*Table S5. Table of all results, regardless of significance*

*Correlation and significance values for all combinations of gene and electrophysiological or morphological feature*

*Table S6. Cell type averages used for analysis of electrophysiological properties*

*Each row represents either an electrophysiological property or a gene. Each column represents one of the 48 cell types defined for the purposes of this analysis, named as “Cre line___layer___cell class.” Each cell contains the mean value of the electrophysiological property, or mean expression level of the gene, within the indicated cell type.*

*Table S7. Cell type averages used for analysis of morphological properties*

*Each row represents either a morphological property or a gene. Each column represents one of the 43 cell types defined for the purposes of this analysis, named as “Cre line___layer___cell class.” Each cell contains the mean value of the morphological property, or mean expression level of the gene, within the indicated cell type.*

## References

1. Gouwens NW, Sorensen SA, Berg J, Lee C, Jarsky T, Ting J, et al. Classification of electrophysiological and morphological types in mouse visual cortex. 2018. Preprint. Available from: https://www.biorxiv.org/content/biorxiv/early/2018/07/17/368456.full.pdf

2. Padmanabhan K, Urban NN. Intrinsic biophysical diversity decorrelates neuronal firing while increasing information content. Nat Neurosci. 2010;13(10):1276–82. Available from: https://doi.org/10.1038/nn.2630

3. Markram H, Muller E, Ramaswamy S, Reimann MW, Defelipe J, Hill SL, et al. Reconstruction and Simulation of Neocortical Microcircuitry. Cell. 2015;163:456–92. Available from: https://doi.org/10.1016/j.cell.2015.09.029

4. Connors BW, Regehr WG. Neuronal firing: Does function follow form? Curr Biol. 1996;6(12):1560–2. Available from: https://doi.org/10.1016/S0960-9822(02)70771-9

5. Chklovskii DB. Synaptic connectivity and neuronal morphology: Two sides of the same coin. Neuron. 2004;43(5):609–17. Available from: https://doi.org/10.1016/j.neuron.2004.08.012

6. Stiefel KM, Sejnowski TJ. Mapping Function Onto Neuronal Morphology. J Neurophysiol. 2007;98(1):513–26. Available from: https://doi.org/10.1152/jn.00865.2006

7. Kopp-Scheinpflug C, Fuchs K, Lippe WR, Tempel BL, Ru R. Decreased Temporal Precision of Auditory Signaling in Kcna1-Null Mice : An Electrophysiological Study In Vivo. J Neurosci. 2003;23(27):9199–207. Available from: https://doi.org/10.1523/JNEUROSCI.23-27-09199.2003

8. Marcotti W, Corns LF, Goodyear RJ, Rzadzinska AK, Avraham KB, Steel KP, et al. The acquisition of mechano-electrical transducer current adaptation in auditory hair cells requires myosin VI. J Physiol. 2016;13:3667–81. Available from: https://doi.org/10.1113/JP272220

9. Qin X, Jiang Y, Tse YC, Wang Y, Wong TP, Paudel HK. Early growth response 1 (Egr-1) regulates N-methyl-D-aspartate receptor (NMDAR)-dependent transcription of PSD-95 and alpha-amino-3-hydroxy-5-methyl-4-isoxazole propionic acid receptor (AMPAR) trafficking in hippocampal primary neurons. J Biol Chem. 2015;290(49):29603–16. Available from: https://doi.org/10.1074/jbc.M115.668889

10. Santiago C, Bashaw GJ. Transcription factors and effectors that regulate neuronal morphology. Development. 2014;141(24):4667–80. Available from: https://doi.org/10.1242/dev.110817

11. Tripathy SJ, Toker L, Li B, Crichlow C, Tebaykin D, Mancarci BO, et al. Transcriptomic correlates of neuron electrophysiological diversity. PLoS Comput Biol. 2017;13(10):1–28. Available from: https://doi.org/10.1371/journal.pcbi.1005814

12. Zeisel A, Muñoz-Manchado AB, Codeluppi S, Lönnerberg P, Manno G La, Juréus A, et al. Cell types in the mouse cortex and hippocampus revealed by single-cell RNA-seq. Science. 2015;347(6226):1138–42. Available from: https://doi.org/10.1126/science.aaa1934

13. Zeisel A, Hochgerner H, Ernfors P, Lo P, Marklund U, Linnarsson S, et al. Molecular Architecture of the Mouse Nervous System. Cell. 2018;174:999–1014. Available from: https://doi.org/10.1016/j.cell.2018.06.021

14. Tasic B, Yao Z, Smith KA, Graybuck L, Nguyen TN, Bertagnolli D, et al. Shared and distinct transcriptomic cell types across neocortical areas. Nature. 2018;563:72–8. Available from: https://doi.org/10.1038/s41586-018-0654-5

15. Tasic B, Menon V, Nguyen TN, Kim TK, Jarsky T, Yao Z, et al. Adult mouse cortical cell taxonomy revealed by single cell transcriptomics. Nat Neurosci. 2016;19(2):335–46. Available from: https://doi.org/10.1038/nn.4216

16. Petilla Interneuron Nomenclature Group, Ascoli GA, Alonso-Nanclares L, Anderson SA, Barrionuevo G, Benavides-Piccione R, et al. Petilla terminology: nomenclature of features of GABAergic interneurons of the cerebral cortex. Nat Rev Neurosci. 2008 Jul 1;9:557. Available from: https://doi.org/10.1038/nrn2402

17. Cadwell CR, Palasantza A, Jiang X, Berens P, Deng Q, Yilmaz M, et al. Electrophysiological, transcriptomic and morphologic profiling of single neurons using Patch-seq. Nat Biotechnol. 2016;34(2):199–203. Available from: https://doi.org/10.1038/nbt.3445

18. Herscovics A. Structure and function of Class I α1,2-mannosidases involved in glycoprotein synthesis and endoplasmic reticulum quality control. Biochimie. 2001;83(8):757–62. Available from: https://doi.org/10.1016/S0300-9084(01)01319-0

19. Huang ZG, Liu HW, Yan ZZ, Wang S, Wang LY, Ding JP. The glycosylation of the extracellular loop of β2 subunits diversifies functional phenotypes of BK Channels. Channels. 2017;11(2):156–66. Available from: https://doi.org/10.1080/19336950.2016.1243631

20. Vicente PC, Kim JY, Ha JJ, Song MY, Lee HK, Kim DH, et al. Identification and characterization of site-specific N-glycosylation in the potassium channel Kv3.1b. J Cell Physiol. 2018;233(1):549–58. Available from: https://doi.org/10.1002/jcp.25915

21. Tripathy SJ, Toker L, Bomkamp C, Mancarci BO, Belmadani M, Pavlidis P. Assessing Transcriptome Quality in Patch-Seq Datasets. Front Mol Neurosci. 2018;11:363. Available from: https://doi.org/10.3389/fnmol.2018.00363

22. Muñoz-Manchado AB, Bengtsson Gonzales C, Zeisel A, Munguba H, Bekkouche B, Skene NG, et al. Diversity of Interneurons in the Dorsal Striatum Revealed by Single-Cell RNA Sequencing and PatchSeq. Cell Rep. 2018;24(8):2179–90. Available from: https://doi.org/10.1016/j.celrep.2018.07.053

23. Brew HM, Hallows JL, Tempel BL. Hyperexcitability reduced low threshold potassium currents auditory neurons of mice lacking the channel subunit Kv1.1. J Physiol. 2003;548(1):1–20.

24. Delprat B, Schaer D, Roy S, Wang J, Puel J-L, Geering K. FXYD6 Is a Novel Regulator of Na,K-ATPase Expressed in the Inner Ear. J Biol Chem. 2007;282(10):7450–6. Available from: https://doi.org/10.1074/jbc.M609872200

25. Harris KD, Hochgerner H, Skene NG, Magno L, Katona L, Bengtsson Gonzales C, et al. Classes and continua of hippocampal CA1 inhibitory neurons revealed by single-cell transcriptomics. PLoS Biol. 2018;16(6):e2006387. Available from: https://doi.org/10.1371/journal.pbio.2006387

26. Fuzik J, Zeisel A, Máté Z, Calvigioni D, Yanagawa Y, Szabó G, et al. Integration of electrophysiological recordings with single-cell RNA-seq data identifies neuronal subtypes. Nat Biotechnol. 2016;34(2):175–83. Available from: https://doi.org/10.1038/nbt.3443

27. Földy C, Darmanis S, Aoto J, Malenka RC, Quake SR, Südhof TC. Single-cell RNAseq reveals cell adhesion molecule profiles in electrophysiologically defined neurons. Proc Natl Acad Sci. 2016;113(35):E5222–31. Available from: https://doi.org/10.1073/pnas.1610155113

28. Islam S, Kjällquist U, Moliner A, Zajac P, Fan JB, Lönnerberg P, et al. Highly multiplexed and strand-specific single-cell RNA 5′ end sequencing. Nat Protoc. 2012;7(5):813–28. Available from: https://doi.org/10.1038/nprot.2012.022

29. Zhu YY, Machleder EM, Chenchik A, Li R, Siebert PD. Reverse transcriptase template switching: A SMART™ approach for full-length cDNA library construction. Biotechniques. 2001;30(4):892–26. 7. Available from: https://doi.org/10.2144/01304pf02

30. Hodgkin AL, Huxley AF. Currents carried by sodium and potassium ions through the membrane of the giant axon of Loligo. J Physiol. 1952;116(4):449–72. Available from: https://doi.org/10.1113/jphysiol.1952.sp004717

31. Stuhmer W, Stocker M, Sakmann B, Seeburg P, Baumann A, Grupe A, et al. Potassium channels expressed from rat brain cDNA have delayed rectifier properties. FEBS Lett. 1988;242(1):199–206. Available from: https://doi.org/10.1016/0014-5793(88)81015-9

32. Alexander SPH, Striessnig J, Kelly E, Marrion N V., Peters JA, Faccenda E, et al. The concise guide to pharmacology 2017/18: Voltage-gated ion channels. Br J Pharmacol. 2017;174:S160–94. Available from: https://doi.org/10.1111/bph.13884

33. Marionneau C, Carrasquillo Y, Norris AJ, Townsend RR, Isom LL, Link AJ, et al. The Sodium Channel Accessory Subunit Nav 1 Regulates Neuronal Excitability through Modulation of Repolarizing Voltage-Gated K+ Channels. J Neurosci. 2012;32(17):5716–27. Available from: https://doi.org/10.1523/JNEUROSCI.6450-11.2012

34. Paisán-Ruiz C. LRRK2 gene variation and its contribution to Parkinson disease. Hum Mutat. 2009;30(8):1153–60. Available from: https://doi.org/10.1002/humu.21038

35. MacLeod D, Dowman J, Hammond R, Leete T, Inoue K, Abeliovich A. The Familial Parkinsonism Gene LRRK2 Regulates Neurite Process Morphology. Neuron. 2006;52(4):587–93. Available from: https://doi.org/10.1016/j.neuron.2006.10.008

36. Dächsel JC, Behrouz B, Yue M, Beevers JE, Melrose HL, Farrer MJ. A comparative study of Lrrk2 function in primary neuronal cultures Justus. Park Relat Disord. 2010;16(10):650–5. Available from: https://doi.org/10.1016/j.parkreldis.2010.08.018

37. Häbig K, Gellhaar S, Heim B, Djuric V, Giesert F, Wurst W, et al. LRRK2 guides the actin cytoskeleton at growth cones together with ARHGEF7 and Tropomyosin 4. Biochim Biophys Acta - Mol Basis Dis. 2013;1832(12):2352–67. Available from: https://doi.org/10.1016/j.bbadis.2013.09.009

38. Borgs L, Peyre E, Alix P, Hanon K, Grobarczyk B, Godin JD, et al. Dopaminergic neurons differentiating from LRRK2 G2019S induced pluripotent stem cells show early neuritic branching defects. Sci Rep. 2016;6:33377. Available from: https://doi.org/10.1038/srep33377

39. Barnat M, Le Friec J, Benstaali C, Humbert S. Huntingtin-Mediated Multipolar-Bipolar Transition of Newborn Cortical Neurons Is Critical for Their Postnatal Neuronal Morphology. Neuron. 2017;93(1):99–114. Available from: https://doi.org/10.1016/j.neuron.2016.11.035

40. Bean BP. The action potential in mammalian central neurons. Nat Rev Neurosci. 2007;8:451–65. Available from: https://doi.org/10.1038/nrn2148

41. Tsuboi T, Fukuda M. Rab3A and Rab27A cooperatively regulate the docking step of dense-core vesicle exocytosis in PC12 cells. J Cell Sci. 2006;119(11):2196–203. Available from: https://doi.org/10.1242/jcs.02962

42. Nakazawa H, Sada T, Toriyama M, Tago K, Sugiura T, Fukuda M, et al. Rab33a Mediates Anterograde Vesicular Transport for Membrane Exocytosis and Axon Outgrowth. J Neurosci. 2012;32(37):12712–25. Available from: https://doi.org/10.1523/JNEUROSCI.0989-12.2012

43. Kelleher RJ, Flanagan PM, Kornberg RD. A Novel Mediator Between Activator Proteins and the RNA Polymerase II Transcription Apparatus. Cell. 1990;61:1209–15. Available from: https://doi.org/10.1016/0092-8674(90)90685-8

44. Ryu S, Zhou S, Ladurner AG, Tjian R. The transcriptional cofactor complex CRSP is required for activity of the enhancer-binding protein Sp1. Nature. 1999;397(6718):446–50. Available from: https://doi.org/10.1038/17141

45. Yin J, Liang Y, Park JY, Chen D, Yao X, Xiao Q, et al. Mediator MED23 plays opposing roles in directing smooth muscle cell and adipocyte differentiation. Genes Dev. 2012;26(19):2192–205. Available from: https://doi.org/10.1101/gad.192666.112

46. Zhu W, Yao X, Liang Y, Liang D, Song L, Jing N, et al. Mediator Med23 deficiency enhances neural differentiation of murine embryonic stem cells through modulating BMP signaling. Development. 2015;142(3):465–76. Available from: https://doi.org/10.1242/dev.112946

47. Hashimoto S, Boissel S, Zarhrate M, Rio M, Munnich A, Egly J, et al. MED23 Mutation Links Intellectual Disability to Dysregulation of Immediate Early Gene Expression. Science. 2011;333:1161–4. Available from: https://doi.org/10.1126/science.1206638

48. Trehan A, Brady JM, Maduro V, Bone WP, Huang Y, Golas GA, et al. MED23-associated intellectual disability in a non-consanguineous family. Am J Med Genet Part A. 2015;167(6):1374–80. Available from: https://doi.org/10.1002/ajmg.a.37047

49. Hoefele J, Sudbrak R, Reinhardt R, Lehrack S, Hennig S, Imm A, et al. Mutational analysis of the NPHP4 gene in 250 patients with nephronophthisis. Hum Mutat. 2005;25(4):411. Available from: https://doi.org/10.1002/humu.9326

50. Yasunaga T, Hoff S, Schell C, Helmstädter M, Kretz O, Kuechlin S, et al. The polarity protein inturned links NPHP4 to daam1 to control the subapical actin network in multiciliated cells. J Cell Biol. 2015;211(5):963–73. Available from: https://doi.org/10.1083/jcb.201502043

51. Xu K, Zhong G, Zhuang X. Actin, Spectrin, and Associated Proteins Form a Periodic Cytoskeletal Structure in Axons. Science. 2013;339:452–6. Available from: https://doi.org/10.1126/science.1232251

52. Ashburner M, Ball CA, Blake JA, Botstein D, Butler H, Cherry JM, et al. Gene ontology: Tool for the unification of biology. Nat Genet. 2000;25(1):25–9. Available from: https://doi.org/10.1038/75556

53. Carbon S, Dietze H, Lewis SE, Mungall CJ, Munoz-Torres MC, Basu S, et al. Expansion of the gene ontology knowledgebase and resources: The gene ontology consortium. Nucleic Acids Res. 2017;45(D1):D331–8. Available from: https://doi.org/10.1093/nar/gkw1108

54. Mancarci BO, Toker L, Tripathy SJ, Li B, Rocco B, Sibille E, et al. Cross-Laboratory Analysis of Brain Cell Type Transcriptomes with Applications to Interpretation of Bulk Tissue Data. Eneuro. 2017;4(6):e0212–17.2017. Available from: https://doi.org/10.1523/ENEURO.0212-17.2017

55. Kluyver T, Ragan-Kelley B, Pérez F, Granger B, Bussonnier M, Frederic J, et al. Jupyter Notebooks — a publishing format for reproducible computational workflows. In: Proceedings of the 20th International Conference on Electronic Publishing. 2016. p. 87–90. Available from: https://doi.org/10.3233/978-1-61499-649-1-87

56. Müllner D. Modern hierarchical, agglomerative clustering algorithms. 2011. Preprint. Available from: http://arxiv.org/abs/1109.2378

57. Bar-Joseph Z, Gifford DK, Jaakkola TS. Fast optimal leaf ordering for hierarchical clustering. Bioinformatics. 2001;17(SUPPL. 1):22–9. Available from: https://doi.org/10.1093/bioinformatics/17.suppl_1.S22

58. Smedley D, Haider S, Durinck S, Pandini L, Provero P, Allen J, et al. The BioMart community portal: An innovative alternative to large, centralized data repositories. Nucleic Acids Res. 2015;43(W1):W589–98. Available from: https://doi.org/10.1093/nar/gkv350

59. Lun ATL, McCarthy DJ, Marioni JC. A step-by-step workflow for low-level analysis of singlecell RNA-seq data with Bioconductor. F1000Research. 2016;5:2122. Available from: https://doi.org/10.12688/f1000research.9501.2

